# Neighbor GWAS: incorporating neighbor genotypic identity into genome-wide association studies of field herbivory

**DOI:** 10.1101/845735

**Authors:** Yasuhiro Sato, Eiji Yamamoto, Kentaro K. Shimizu, Atsushi J. Nagano

## Abstract

An increasing number of field studies have shown that the phenotype of an individual plant depends not only on its genotype but also on those of neighboring plants; however, this fact is not taken into consideration in genome-wide association studies (GWAS). Based on the Ising model of ferromagnetism, we incorporated neighbor genotypic identity into a regression model, named “Neighbor GWAS”. Our simulations showed that the effective range of neighbor effects could be estimated using an observed phenotype from when the proportion of phenotypic variation explained (PVE) by neighbor effects peaked. The spatial scale of the first nearest neighbors gave the maximum power to detect the causal variants responsible for neighbor effects, unless their effective range was too broad. However, if the effective range of the neighbor effects was broad and minor allele frequencies were low, there was collinearity between the self and neighbor effects. To suppress the false positive detection of neighbor effects, the fixed effect and variance components involved in the neighbor effects should be tested in comparison with a standard GWAS model. We applied neighbor GWAS to field herbivory data from 199 accessions of *Arabidopsis thaliana* and found that neighbor effects explained 8% more of the PVE of the observed damage than standard GWAS. The neighbor GWAS method provides a novel tool that could facilitate the analysis of complex traits in spatially structured environments and is available as an R package at CRAN (https://cran.rproject.org/package=rNeighborGWAS).

## INTRODUCTION

Plants are immobile and thus cannot escape their neighbors. In natural and agricultural systems, individual phenotypes depend not only on the plants’ own genotype but also on the genotypes of other neighboring plants (Tahvanainen and Root 1972; Barbosa et al. 2009; Underwood et al. 2014). This phenomenon has been termed neighbor effects or associational effects in plant ecology (Barbosa et al. 2009; Underwood et al. 2014; Sato 2018). Such neighbor effects were initially reported as a form of interspecific interaction among different plant species (Tahvanainen and Root 1972), but many studies have illustrated that neighbor effects occur among different genotypes within a plant species with respect to: (i) herbivory (Schuman et al. 2015; Sato 2018; Ida et al. 2018), (ii) pathogen infections (Mundt 2002; Zeller et al. 2012), and (iii) pollinator visitations (Underwood et al. 2020). Although neighbor effects are of considerable interest in plant science (Dicke and Baldwin 2010; Erb 2018) and agriculture (Zeller et al. 2012; Dahlin et al. 2018), they are often not considered in quantitative genetic analyses of field-grown plants.

Complex mechanisms underlie neighbor effects through direct competition (Weiner 1990), herbivore and pollinator movement (Bergvall et al. 2006; Verschut et al. 2016; Underwood et al. 2020), and volatile communication among plants (Schuman et al. 2015; Dahlin et al. 2018). For example, lipoxygenase (*LOX)* genes govern jasmonate-mediated volatile emissions in wild tobacco (*Nicotiana attenuata)* that induce defenses of neighboring plants (Schuman et al. 2015). Even if direct plant–plant communications are absent, herbivores can mediate indirect interactions between plant genotypes (Sato and Kudoh 2017; Ida et al. 2018). For example, the *GLABRA1* gene is known to determine hairy or glabrous phenotypes in *Arabidopsis* plants (Hauser et al. 2001), and the flightless leaf beetle (*Phaedon brassicae)* is known to prefer glabrous plants to hairy ones (Sato et al. 2017). Consequently, hairy plants escape herbivory when surrounded by glabrous plants (Sato and Kudoh 2017). Yet, there are few hypothesis-free approaches currently available for the identification of the key genetic variants responsible for plant neighborhood effects.

Genome-wide association studies (GWAS) have been increasingly adopted to resolve the genetic architecture of complex traits in the model plant, *Arabidopsis thaliana* (Atwell et al. 2010; Seren et al. 2017; Togninalli et al. 2018), and crop species (Hamblin et al. 2011). The interactions of plants with herbivores (Brachi et al. 2015; Nallu et al. 2018), microbes (Horton et al. 2014; Wang et al. 2018), and other plant species (Frachon et al. 2019) are examples of the complex traits that are investigated through the lens of GWAS. To distinguish causal variants from the genome structure, GWAS often employs a linear mixed model with kinship considered as a random effect (Kang et al. 2008; Korte and Farlow 2013). However, because of combinatorial explosion, it is generally impossible to test the full set of inter-genomic locus-by-locus interactions (Gondro et al. 2013); thus, some feasible and reasonable approach should be developed for the GWAS of neighbor effects.

To incorporate neighbor effects into GWAS, we have focused on a theoretical model of neighbor effects in magnetic fields, known as the Ising model (Ising 1925; McCoy and Maillard 2012), which has been applied to forest gap dynamics (Kizaki and Katori 1999; Schlicht and Iwasa 2004) and community assembly (Azaele et al. 2010) in plant ecology. Using the Ising analogy, we compare individual plants to a magnet: the two alleles at each locus correspond to the north and south dipoles, and genome-wide multiple testing across all loci is analogous to a number of parallel two-dimensional layers. The Ising model has a clear advantage in its interpretability, such that: (i) the optimization problem for a population sum of trait values can be regarded as an inverse problem of a simple linear model, (ii) the sign of neighbor effects determines the model’s trend with regard to the generation of a clustered or checkered spatial pattern of the two states, and (iii) the self-genotypic effect determines the general tendency to favor one allele over another (Fig. 1).

**Figure 1.**
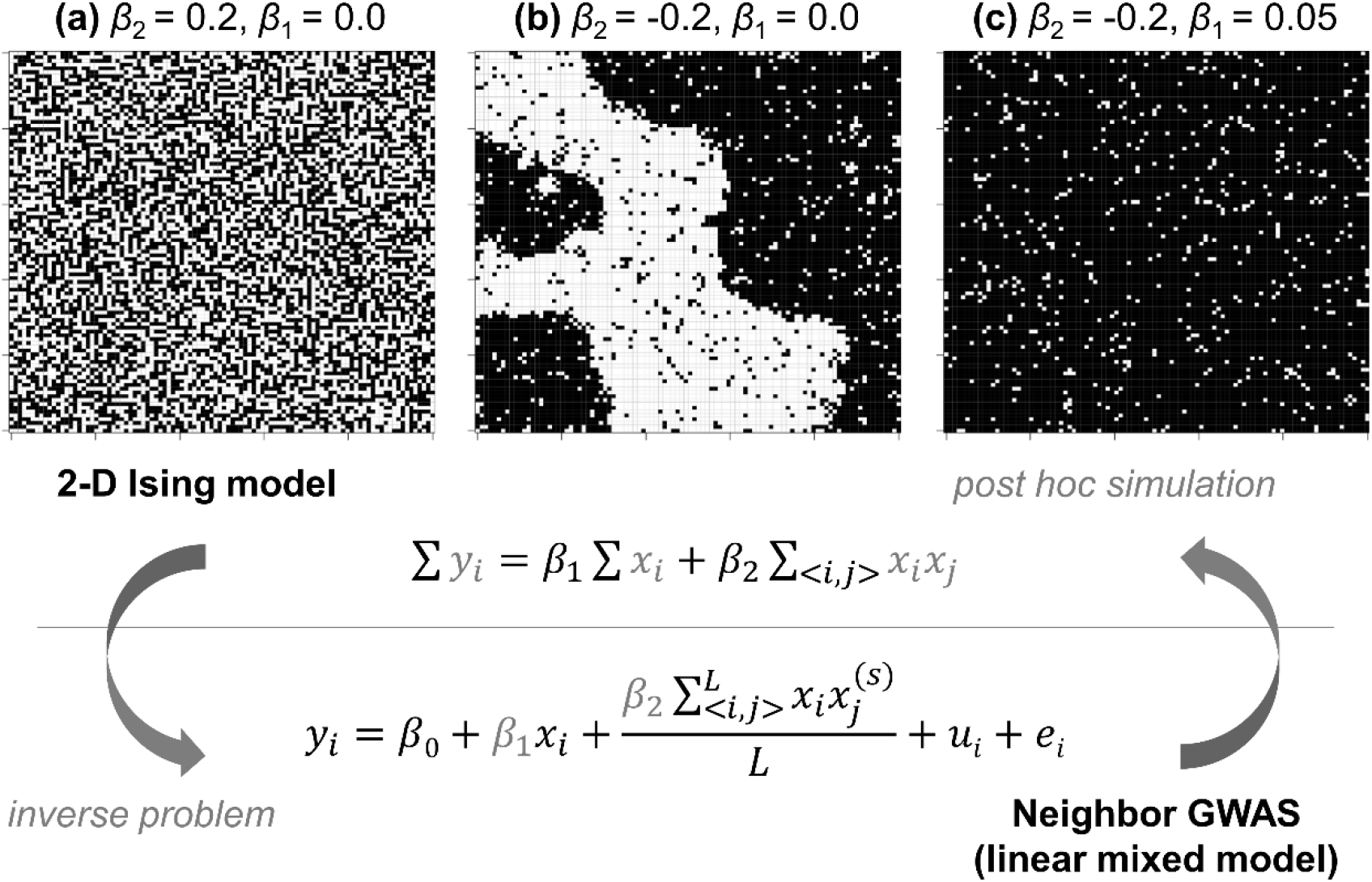
Relationship between the neighbor GWAS and Ising model. Upper panels show the spatial arrangements expected by a 2-D Ising model ∑*y_i_* = *β*_1_∑*x_i_* + *β*_2_ ∑_<*i,j*_> *x_i_x_j_*. (a) If *β*_2_>0, mixed patterns give the argument of the minimum for a population sum of phenotype values ∑*y_i_*. (b) If *β*_2_<0, clustered patterns give the argument of the minimum for ∑*y_i_*. (c) In addition, *β*_1_ determines the overall patterns favoring −1 or +1 states. The figures show outcomes from a random 100 × 100 lattice after 1000 iterations of simulated annealing. Conversely, the neighbor GWAS was implemented as an inverse problem of the 2-D Ising model, where genotypes and its spatial arrangement, *x_i_* and *x_i_x_j_*, were given while the coefficients *β*_1_ and *β*_2_ were to be estimated from the observed phenotypes *y_i_*. In addition, the variance component due to self and neighbor effects was considered a random effect in a linear mixed model, such that *u_i_* ∈ ***u*** and 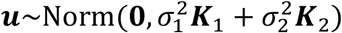. Once *β*_1_ and *β*_2_ were determined, we could simulate a genotype distribution that maximizes or minimizes ∑*y_i_*.

In this study, we proposed a new methodology integrating GWAS and the Ising model, named “neighbor GWAS.” The method was applied to simulated phenotypes and actual data of field herbivory on *A. thaliana.* We addressed two specific questions: (i) what spatial and genetic factors influenced the power to detect causal variants? and (ii) were neighbor effects significant sources of leaf damage variation in field-grown *A. thaliana*? Based on the simulation and application, we determined the feasibility of our approach to detect neighbor effects in field-grown plants.

## MATERIALS & METHODS

### Neighbor GWAS

#### Basic model

We analyzed neighbor effects in GWAS as an inverse problem of the two-dimensional Ising model, named “neighbor GWAS” hereafter (Fig. 1). We considered a situation where a plant accession has one of two alleles at each locus, and a number of accessions occupied a finite set of field sites, in a two-dimensional lattice. The allelic status at each locus was represented by *x*, and so the allelic status at each locus of the *i*-th focal plant and the *j*-th neighboring plants was designated as *x*_*i*(*j*)_ ∈{-1, +1}. Based on a two-dimensional Ising model, we defined a phenotype value for the *i*-th focal individual plant *y_i_* as:

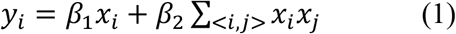

where *β*_1_ and *β*_2_ denoted self-genotype and neighbor effects, respectively. If two neighboring plants shared the same allele at a given locus, the product *x_i_x_j_* turned into (−1) × (−1) = +1 or (+1) × (+1) = +1. If two neighbors had different alleles, the product *x_i_x_j_* became (−1) × (+1) = −1 or (+1) × (−1) = −1. Accordingly, the effects of neighbor genotypic identity on a particular phenotype depended on the coefficient *β*_2_ and the number of the two alleles in a neighborhood. If the numbers of identical and different alleles were the same near a focal plant, these neighbors offset the sum of the products between the focal plant *i* and all *j* neighbors ∑_<*i,j*>_ *x_i_x_j_* and exerted no effects on a phenotype. When we summed up the phenotype values for the total number of plants *n* and replaced it as *E* = −*β*_2_, *H* = −*β*_1_ and *ϵ_I_* = ∑*y_i_*, eq. 1 could be transformed into *ϵ_I_* = −*E*∑_<*i,j*>_ *x_i_x_j_* – *H*∑*x_i_*, which defined the interaction energy of a two-dimensional ferromagnetic Ising model (McCoy and Maillard 2012). The neighbor effect *β*_2_ and self-genotype effect *β*_1_ were interpreted as the energy coefficient *E* and external magnetic effects *H*, respectively. An individual plant represented a spin and the two allelic states of each locus corresponded to a north or south dipole. The positive or negative value of ∑*x_i_x_j_* indicated a ferromagnetism or paramagnetism, respectively. In this study, we did not consider the effects of allele dominance because this model was applied to inbred plants. However, heterozygotes could be processed if the neighbor covariate *x_i_x_j_* was weighted by an estimated degree of dominance in the self-genotypic effects on a phenotype.

#### Association tests

For association mapping, we needed to determine *β*_1_ and *β*_2_ from the observed phenotypes and considered a confounding sample structure as advocated by previous GWAS (e.g., Kang et al. 2008; Korte and Farlow 2013). Extending the basic model (eq. 1), we described a linear mixed model at an individual level as:

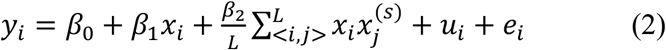

where *β*_0_ indicated the intercept, and the term *β*_1_*x_i_* represented fixed self-genotype effects as tested in standard GWAS; *β*_2_ was the coefficient of fixed neighbor effects. The neighbor covariate 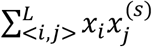 indicated a sum of products for all combinations between the *i*-th focal plant and the *j*-th neighbor at the *s*-th spatial scale from the focal plant *i*, and was scaled by the number of neighboring plants, *L.* The number of neighboring plants *L* was dependent on the spatial scale s to be referred. Variance components due to the sample structure of self and neighbor effects were modeled by a random effect *u_i_*, ∈ ***u*** and 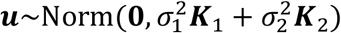. The residual was expressed as *e_i_* ∈ ***e*** and 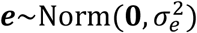.

#### Variation partitioning

To estimate the proportion of phenotypic variation explained (PVE) by the self and neighbor effects, we utilized variance component parameters in linear mixed models. The *n* × *n* variance-covariance matrices represented the similarity in self-genotypes (i.e., kinship) and neighbor covariates among *n* individual plants as 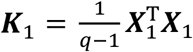 and 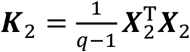, where *q* indicated the number of markers. As we defined *x*_*i*(*j*)_ ∈{+1, −1}, the elements of the kinship matrix ***K***_1_ were scaled to represent the proportion of marker loci shared among *n* × *n* plants such that 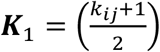; 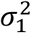 and 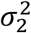 indicated variance component parameters for the self and neighbor effects.

The elements of *n* plants × *q* markers matrix ***X***_1_ and ***X***_2_ consisted of explanatory variables for the self and neighbor effects as ***X***_1_ = (*x_i_*) and 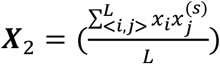. The individual-level formula eq. 2 could also be converted into a conventional matrix form as:

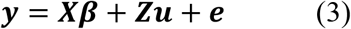

where ***y*** was an *n* × 1 vector of the phenotypes; ***X*** was a matrix of fixed effects, including a unit vector, self-genotype *x_i_*, neighbor covariate 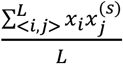, and other confounding covariates for *n* plants; ***β*** was a vector that represents the coefficients of the fixed effects; ***Z*** was a design matrix allocating individuals to a genotype, and became an identity matrix if all plants were different accessions; ***u*** was the random effect with 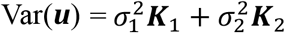; and ***e*** was residual as 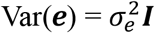.

Because our objective was to test for neighbor effects, we needed to avoid the detection of false positive neighbor effects. The self-genotype value *x_i_* and neighbor genotypic identity 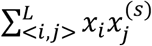 would be colinear due to the minor allele frequency (MAF) and the spatial scale of *s*. When MAF is low, neighbors 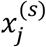 are unlikely to vary in space and most plants will have similar values for neighbor identity 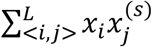. Furthermore, if the neighbor effects range was broad enough to encompass an entire field (i.e., *s* → ∞), the neighbor covariate and self-genotype *x_i_* would become colinear according to the equation: 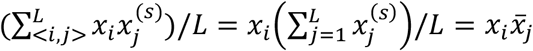, where 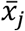 indicates a population-mean of neighbor genotypes and corresponds to a population-mean of self-genotype values 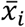, if *s* → ∞. The standard GWAS is a subset of the neighbor GWAS and these two models become equivalent at *s* = 0 and 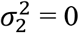. When testing the self-genotype effect *β*_1_, we recommend that the neighbor effects and its variance component 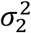 should be excluded; otherwise, the standard GWAS fails to correct a sample structure because of the additional variance component at 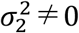. To obtain a conservative conclusion, the significance of *β*_2_ and 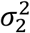 should be compared using the standard GWAS model based on self-effects alone.

Given the potential collinearity between the self and neighbor effects, we defined different metrics for the proportion of phenotypic variation explained (PVE) based on self or neighbor effects. Using a single-random effect model, we calculated PVE for either the self or neighbor effects as follows:

‘single’ 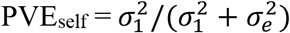 when *s* and 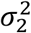 were set at 0, or ‘single’ 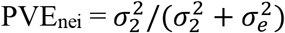 when 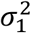 was set at 0.

Furthermore, we could partial out either of the two variance components using a two-random effect model and define PVE as:

‘partial’ 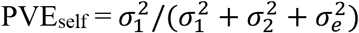 and ‘partial’ 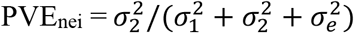.

As the partial PVE_self_ was equivalent to the single PVE_self_ when *s* was set at 0, the net contribution of neighbor effects at *s* ≠ 0 was given as ‘net’ PVE_nei_ = (partial PVE_self_ + partial PVE_nei_) – single PVE_self_, which indicated the proportion of phenotypic variation that could be explained by neighbor effects, but not by the self-genotype effects.

### Simulation

To examine the model performance, we applied the neighbor GWAS to simulated phenotypes. Phenotypes were simulated using a subset of the actual *A. thaliana* genotypes. To evaluate the performance of the simple linear model, we assumed a complex ecological form of neighbor effects with multiple variance components controlled. The model performance was evaluated in terms of the causal variant detection and accuracy of estimates. All analyses were performed using R version 3.6.0 (R Core Team 2019).

#### Genotype data

To consider a realistic genetic structure in the simulation, we used part of the *A. thaliana* RegMap panel (Horton et al. 2012). The genotype data for 1,307 accessions were downloaded from the Joy Bergelson laboratory website (http://bergelson.uchicago.edu/?page_id=790 accessed on February 9, 2017). We extracted data for chromosomes 1 and 2 with MAF at >0.1, yielding a matrix of 1,307 plants with 65,226 single nucleotide polymorphisms (SNPs). Pairwise linkage disequilibrium (LD) among the loci was *r*^2^ = 0.003 [0.00-0.06: median with upper and lower 95 percentiles]. Before generating a phenotype, genotype values at each locus were standardized to a mean of zero and a variance of 1. Subsequently, we randomly selected 1,296 accessions (= 36 × 36 accessions) without any replacements for each iteration and placed them in a 36 × 72 checkered space, following the *Arabidopsis* experimental settings (see Fig. S1).

#### Phenotype simulation

To address ecological issues specific to plant neighborhood effects, we considered two extensions, namely asymmetric neighbor effects and spatial scales. Previous studies have shown that plant–plant interactions between accessions are sometimes asymmetric under herbivory (e.g., Bergvall et al. 2006; Verschut et al. 2016; Sato and Kudoh 2017) and height competition (Weiner 1990); where one focal genotype is influenced by neighboring genotypes, while another receives no neighbor effects. Such asymmetric neighbor effects can be tested by statistical interaction terms in a linear model (Bergvall et al. 2006; Sato and Kudoh 2017). Several studies have also shown that the strength of neighbor effects depends on spatial scales (Hambäck et al. 2014), and that the scale of neighbors to be analyzed relies on the dispersal ability of the causative organisms (see Hambäck et al. 2009; Sato and Kudoh 2015; Verschut et al. 2016; Ida et al. 2018 for insect and mammal herbivores; Rieux et al. 2014 for pathogen dispersal) or the size of the competing plants (Weiner 1990). We assumed the distance decay at the *s*-th sites from a focal individual *i* with the decay coefficient *α* as *w*(*s, α*) = e^−*α*(*s*−1)^, since such an exponential distance decay has been widely adopted in empirical studies (Devaux et al. 2007; Carrasco et al. 2010; Rieux et al. 2014; Ida et al. 2018). Therefore, we assumed a more complex model for simulated phenotypes than the model for neighbor GWAS as follows:

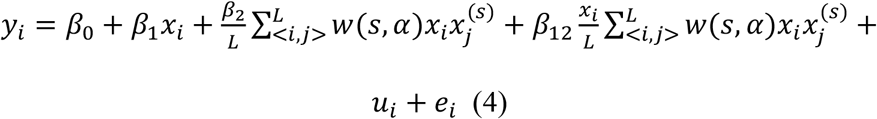

where *β*_12_ was the coefficient for asymmetry in neighbor effects. By incorporating an asymmetry coefficient, the model (eq. 4) can deal with cases where neighbor effects are one-sided or occur irrespective of a focal genotype (Fig. 2). Total variance components resulting from three background effects (i.e., the self, neighbor, and self-by-neighbor effects) were defined as *u_i_* ∈ ***u*** and 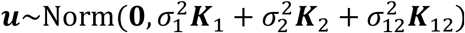. The three variance component parameters 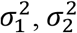, and 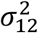, determined the relative importance of the self-genotype, neighbor, and asymmetric neighbor effects in u,. Given the elements of *n* plants × *q* marker explanatory matrix with 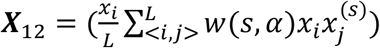, the similarity in asymmetric neighbor effects was calculated as 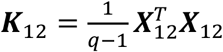. To control phenotypic variations, we further partitioned the proportion of phenotypic variation into those explained by the major-effect genes and variance components PVE_***β***_ + PVE_***u***_, major-effect genes alone PVE_***β***_ and residual error PVE_***e***_, where PVE_***β***_ + PVE_***u***_ + PVE_***e***_ = 1. The *optimize* function in R was used to adjust the simulated phenotypes to the given amounts of PVE.

**Figure 2.**
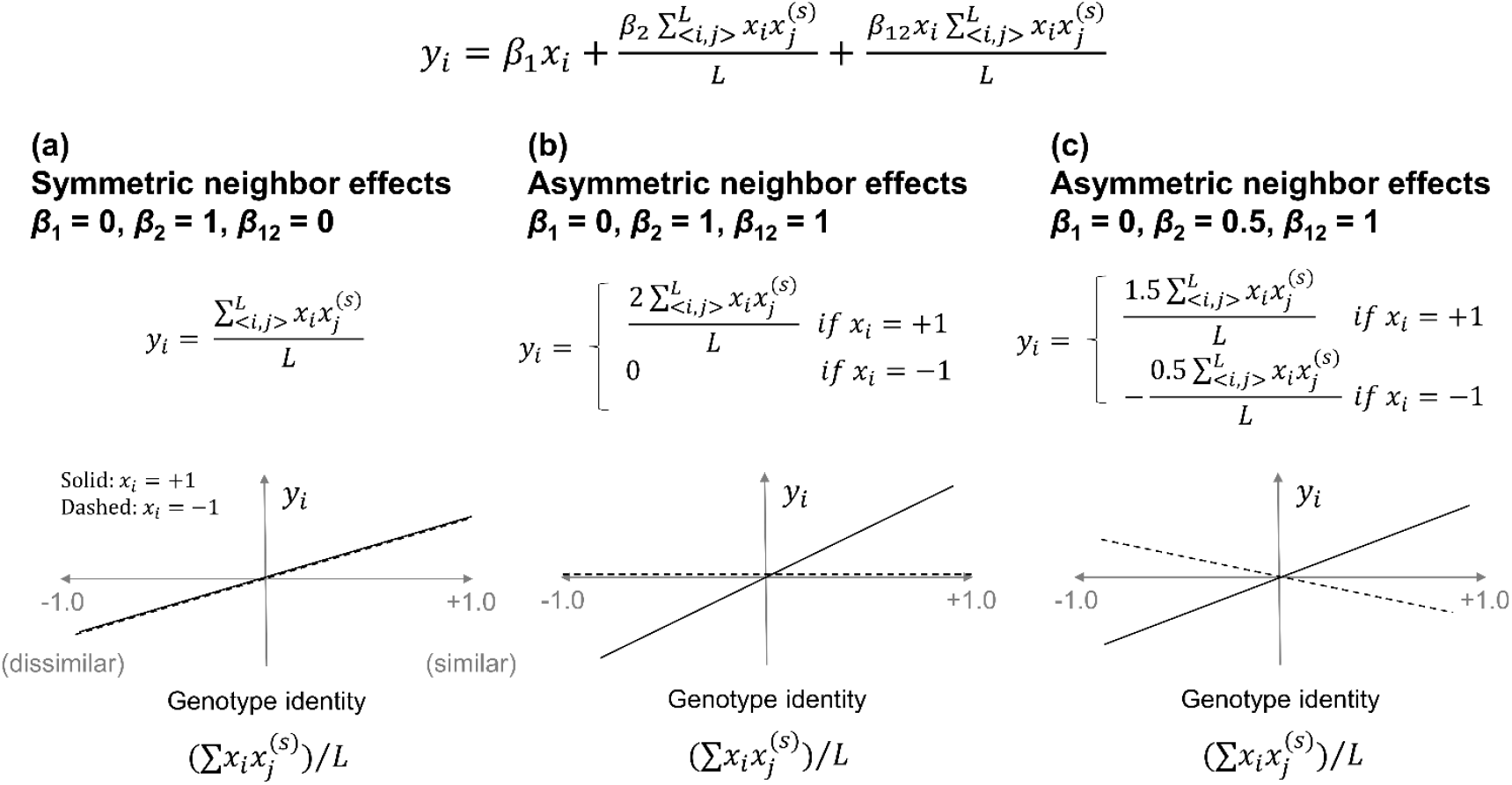
Numerical examples of the symmetric (a) and asymmetric (b, c) neighbor effects. The intercept, distance decay, random effects, and residual errors are neglected, to simplify this scheme. (a) Symmetric neighbor effects represent how neighbor genotype similarity (or dissimilarity) affects the trait value of a focal individual *y_i_* regardless of its own genotype. (b) Asymmetric neighbor effects can represent a case in which one genotype experiences neighbor effects while the other does not (b) and a case in which the direction of the neighbor effects depends on the genotypes of a focal individual (c). The case (b) was considered in our simulation as it has been empirically reported (e.g., Bergvall et al. 2006; Verschut et al. 2016; Sato & Kudoh 2017).

#### Parameter setting

Ten phenotypes were simulated with varying combination of the following parameters, including the distance decay coefficient *α*, the proportion of phenotypic variation explained by the major-effect genes PVE_***β***_ the proportion of phenotypic variation explained by major-effect genes and variance components PVE_***β***_ + PVE_***u***_, and the relative contributions of self, symmetric neighbor, and asymmetric neighbor effects, i.e., PVE_self_:PVE_nei_:PVE_s×n_. We run the simulation with different combinations, including *α* = 0.01, 1.0, or 3.0; PVE_self_:PVE_nei_:PVE_s×n_ = 8:1:1, 5:4:1, or 1:8:1; and PVE_***β***_ and PVH_***β***_ + PVE_***u***_ = 0.1 and 0.4, 0.3 and 0.4, 0.3 and 0.8, or 0.6 and 0.8. The maximum reference scale was fixed at *s* = 3.

The line of simulations was repeated for 10, 50, or 300 causal SNPs to examine cases of oligogenic and polygenic control of a trait. The non-zero coefficients for the causal SNPs were randomly sampled from −1 or 1 digit and then assigned, as some causal SNPs were responsible for both the self and neighbor effects. Of the total number of causal SNPs, 15% had self, neighbor, and asymmetric neighbor effects (i.e., *β*_1_ ≠ 0 and *β*_2_ ≠ 0 and *β*_12_ ≠ 0); another 15% had both the self and neighbor effects, but no asymmetry in the neighbor effects (*β*_1_ ≠ 0 and *β*_2_ ≠ 0 and *β*_12_ = 0); another 35% had self-genotypic effects only (*β*_1_ ≠ 0); and the remaining 35% had neighbor effects alone (*β*_2_ ≠ 0). Given its biological significance, we assumed that some loci having neighbor signals, possessed asymmetric interactions between the neighbors (*β*_2_ ≠ 0 and *β*_12_ ≠ 0), while the others had symmetric interactions (*β*_2_ ≠ 0 and *β*_12_ = 0). Therefore, the number of causal SNPs in *β*_12_ was smaller than that in the main neighbor effects *β*_2_. According to this assumption, the variance component 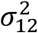 was also assumed to be smaller than 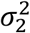. To examine extreme conditions and strong asymmetry in neighbor effects, we additionally analyzed the cases with PVE_self_:PVE_nei_:PVE_s×n_ = 1:0:0, 0:1:0, or 1:1:8.

#### Summary statistics

The simulated phenotypes were fitted by eq. 2 to test the significance of coefficients *β*_1_ and *β*_2_, and to estimate single or partial PVE_self_ and PVE_nei_. To deal with potential collinearity between *x_i_* and neighbor genotypic identity 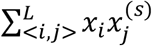, we performed likelihood ratio tests between the self-genotype effect model and the model with both self and neighbor effects, which resulted in conservative tests of significance for *β_2_* and 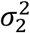. The simulated phenotype values were standardized to have a mean of zero and a variance of 1, where true *β* was expected to match the estimated coefficients 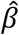 when multiplied by the standard deviation of non-standardized phenotype values. The likelihood ratio was calculated as the difference in deviance, i.e., −2 × log-likelihood, which is asymptotically *χ*^2^ distributed with one degree of freedom. The variance components, 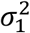 and 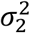, were estimated using a linear mixed model without any fixed effects. To solve the mixed model with the two random effects, we used the average information restricted maximum likelihood (AI-REML) algorithm implemented in the *Imm.aireml* function in the *gaston* package of R (Perdry and Dandine-Roulland 2018). Subsequently, we replaced the two variance parameters 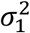 and 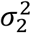 in eq. 2 with their estimates 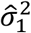 and 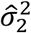 from the AI-REML, and performed association tests by solving a linear mixed model with a fast approximation, using eigenvalue decomposition (implemented in the *Imm.diago* function: Perdry and Dandine-Roulland 2018). The model likelihood was computed using the *Imm.diago.profile.likelihood* function. We evaluated the self and neighbor effects for association mapping based on the forward selection of the two fixed effects, *β*_1_ and *β*_2_, as described below:

1. Computed the null likelihood with 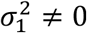 and 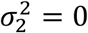 in eq. 2
2. Tested the self-effect, *β*_1_, by comparing with the null likelihood
3. Computed the self-likelihood with 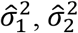 and *β*_1_ using eq. 2

Tested the neighbor effects, *β*_2_, by comparing with the self-likelihood We also calculated PVE using the mixed model (eq. 3) without *β*_1_ and *β*_2_ as follows:

1. Calculated single PVE_self_ or single PVE_nei_ by setting either 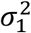 or 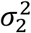 at 0.
2. Tested the single PVE_self_ or single PVE_nei_ using the likelihood ratio between the null and one-random effect model
3. Calculated the partial PVE_self_ and partial PVE_nei_ by estimating 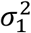 and 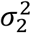 simultaneously
4. Tested the partial PVE_self_ and partial PVE_nei_ using the likelihood ratio between the two- and one-random effect model

We inspected the model performance based on causal variant detection, PVE estimates, and effect size estimates. The true or false positive rates between the causal and non-causal SNPs were evaluated using ROC curves and area under the ROC curves (AUC) (Gage et al. 2018). An AUC of 0.5 would indicate that the GWAS has no power to detect true signals, while an AUC of 1.0 would indicate that all the top signals predicted by the GWAS agree with the true signals. In addition, the sensitivity to detect self or neighbor signals (i.e., either *β*_1_ ≠ 0 or *β*_2_ ≠ 0) was evaluated using the true positive rate of the ROC curves at a stringent specificity level, where the false positive rate = 0.05. The roc function in the pROC package (Robin et al. 2011) was used to calculate the ROC and AUC from −log_10_(*p*-value). Factors affecting the AUC or sensitivity were tested by analysis-of-variance (ANOVA) for the self or neighbor effects (AUC_self_ or AUC_nei_; self or neighbor sensitivity). The AUC and PVE were calculated from *s* = 1 (the first nearest neighbors) to *s* = 3 (up to the third nearest neighbors) cases. The AUC was also calculated using standard linear models without any random effects, to examine whether the linear mixed models were superior to the linear models. We also tested the neighbor GWAS model incorporating the neighbor phenotype 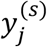. instead of 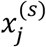. The accuracy of the total PVE estimates was defined as PVE accuracy = (estimated total PVE – true total PVE) / true total PVE. The accuracy of the effect size estimates was evaluated using mean absolute errors (MAE) between the true and estimated (*β*_1_ or *β*_2_ for the self and neighbor effects (MAE_self_ and MAE_nei_). Factors affecting the accuracy of PVE and effect size estimates were also tested using ANOVA. Misclassifications between self and neighbor fixed effects were further evaluated by comparing *p*-value scores between zero and non-zero coefficients. If −log_10_(*p*-value) scores of zero *β* are the same or larger than non-zero *β*, it infers a risk of misspecification of the true signals.

### *Arabidopsis* herbivory data

We applied the neighbor GWAS to field data of *Arabidopsis* herbivory. The procedure for this field experiment followed that of our previous experiment (Sato et al. 2019). We selected 199 worldwide accessions from 2029 accessions sequenced by the RegMap (Horton et al. 2012) and 1001 Genomes project (Alonso-Blanco et al. 2016). Of the 199 accessions, most were overlapped with a previous GWAS of biotic interactions (Horton et al. 2014) and half were included by a GWAS of glucosinolates (Chan et al. 2010). Eight replicates of each of the 199 accessions were first prepared in a laboratory and then transferred to the outdoor garden at the Center for Ecological Research, Kyoto University, Japan (Otsu, Japan: 35°06’N, 134°56?, alt. ca. 200 m: Fig. S1). Seeds were sown on Jiffy-seven pots (33-mm diameter) and stratified at a temperature of 4 ***°C*** for a week. Seedlings were cultivated for 1.5 months under a short-day condition (8 h light: 16 h dark, 20 °C). Plants were then separately potted in plastic pots (6 cm in diameter) filled with mixed soil of agricultural compost (Metro-mix 350, SunGro Co., USA) and perlite at a 3:1 ratio. Potted plants were set in plastic trays (10 × 40 cells) in a checkered pattern (Fig. S1). In the field setting, a set of 199 accessions and an additional Col-0 accession were randomly assigned to each block without replacement (Fig. S1). Eight replicates of these blocks were set >2 m apart from each other (Fig. S1). Potted plants were exposed to the field environment for 3 wk in June 2017. At the end of the experiment, the percentage of foliage eaten was scored as: 0 for no visible damage, 1 for ≤10%, 2 for >10% and ≤ 25%, 3 for > 25% and ≤ 50%, 4 for >50% and ≤ 75%, and 5 for >75%. All plants were scored by a single person to avoid observer bias. The most predominant herbivore in this field trial was the diamond back moth (*Plutella xylostella)*, followed by the small white butterfly (*Pieris rapae).* We also recorded the initial plant size and the presence of inflorescence to incorporate them as covariates. Initial plant size was evaluated by the length of the largest rosette leaf (mm) at the beginning of the field experiment and the presence of inflorescence was recorded 2 wk after transplanting.

We estimated the variance components and performed the association tests for the leaf damage score with the neighbor covariate at *s* = 1 and 2. These two scales corresponded to *L* = 4 (the nearest four neighbors) and *L* = 12 (up to the second nearest neighbors), respectively, in the *Arabidopsis* dataset. The variation partitioning and association tests were performed using the *gaston* package, as mentioned above. To determine the significance of the variance component parameters, we compared the likelihood between mixed models with one or two random effects. For the genotype data, we used an imputed SNP matrix of the 2029 accessions studied by the RegMap (Horton et al. 2012) and 1001 Genomes project (Alonso-Blanco et al. 2016). Missing genotypes were imputed using BEAGLE (Browning and Browning 2009), as described by Togninalli et al. (2018) and updated on the AraGWAS Catalog (https://aragwas.1001genomes.org). Of the 10,709,466 SNPs from the full imputed matrix, we used 1,242,128 SNPs with MAF at > 0.05 and LD of adjacent SNPs at *r*^2^ <0.8. We considered the initial plant size, presence of inflorescence, experimental blocks, and the edge or center within a block as fixed covariates; these factors explained 12.5% of the leaf damage variation (1.2% by initial plant size, Wald test, *Z* = 3.53, *p*-value<0.001; 2.4% by the presence of inflorescence, *Z* = −5.69, *p*-value<10^-8^; 8.3% by the experimental blocks, likelihood ratio test, χ^2^ = 152.8, df = 7, *p*-value<10^-28^; 0.5% by the edge or center, *Z* = 3.11, *p*-value = 0.002). After the association mapping, we searched candidate genes within ~10 kb around the target SNPs, based on the Araport11 gene model with the latest annotation of The Arabidopsis Information Resource (TAIR) (accessed on 7 September 2019). Gene-set enrichment analysis was performed using the Gowinda algorithm that enables unbiased analysis of the GWAS results (Kofler and Schlotterer 2012). We tested the SNPs with the top 0.1% −log_10_(*p*-value) scores, with the option “--gene-definition undownstream10000,” “--min-genes 20,” and “--mode gene.” The GO.db package (Carlson et al. 2018) and the latest TAIR AGI code annotation were used to build input files. The R source codes, accession list, and phenotype data are available at the GitHub repository (https://github.com/naganolab/NeighborGWAS).

### R package, “rNeighborGWAS”

To increase the availability of the new method, we have developed the neighbor GWAS into an R package, which is referred to as “rNeighborGWAS”. In addition to the genotype and phenotype data, the package requires a spatial map indicating the positions of individuals across a space. In this package, we generalized the discrete space example into a continuous two-dimensional space, allowing it to handle any spatial distribution along the x- and y-axes. Based on the three input files, the rNeighborGWAS package estimates the effective range of neighbor effects by calculating partial PVE_nei_ and performs association mapping of the neighbor effects using the linear mixed models described earlier. Details and usage are described in the help files and vignette of the rNeighborGWAS package available via CRAN at https://cran.r-project.org/package=rNeighborGWAS.

To assess its implementation, we performed standard GWAS using GEMMA version 0.98 (Zhou and Stephens 2012) and the rNeighborGWAS. The test phenotype data were the leaf damage scores for the 199 accessions described previously and their flowering times under long-day conditions (“FT16” phenotype collected by Atwell et al. 2010 and Alonso-Blanco et al. 2016). The flowering time phenotype was downloaded from the AraPheno database (https://arapheno.1001genomes.org/: Seren et al. 2017). The full imputed genotype data were compiled for 1057 accessions, whose genotypes and flowering time phenotype were both available. The cut-off value of the MAF was set at 5%, yielding 1,814,755 SNPs for the 1057 accessions. The same kinship matrix defined by ***K***_1_ above was prepared as an input file. We calculated *p*-values using likelihood ratio tests in the GEMMA program, because the rNeighborGWAS adopted likelihood ratio tests.

## RESULTS

### Simulation

We conducted simulations to test the capability of the neighbor GWAS to estimate PVE and marker-effects. As expected by the model and data structure, collinearity was detected between the self-genotypic variable *x_i_* and the neighbor variable ∑*x_i_x_j_*/*L* in the simulated genotypes (Fig. S2). The level of collinearity varied from a slight correlation to complete collinearity as the MAF became smaller, from 0.5 to 0.1 (Fig. S2). The collinearity was also more severe as the scale of *s* was increased. For example, even at *s* = 2, we could cut off the MAF at >0.4 to keep |*r*| below 0.6 for all SNPs. The element-wise correlation between ***K**_1_* and ***K***_2_ indicated that at least 60% of the variation was overlapping between the two genome-wide variance-covariance matrices in the partial genotype data used for this simulation (*R*^2^ = 0.62 at *s* = 1; *R*^2^ = 0.79 at *s* = 2; *R*^2^ = 0.84 at *s* = 3).

A set of phenotypes were then simulated from the real genotype data following a complex model (eq. 4), and then fitted using a simplified model (eq. 2). The accuracy of the total PVE estimation was the most significantly affected by the spatial scales of *s* (Table 1). The total PVE was explained relatively well by the single PVE_self_ that represented the additive polygenic effects of the self-genotypes (Fig. 3). Inclusion of partial PVE_nei_ accounted for the rest of the true total PVE, which was considered the net contribution of neighbor effects to phenotypic variation. The net PVE_nei_ was largest when the effective range of the neighbor effects was narrow (i.e., strong distance decay at *α* = 3) and the contribution of the partial PVE_nei_ was much larger than that of PVE_self_ (Fig. 3). However, the sum of the single PVE_self_ (= partial PVE_self_ at *s* = 0) and the partial PVE_nei_ did not match the true total PVE (Fig. 3), as expected by the collinearity between the self and neighbor effects (Fig. S2). Due to such collinearity, the single PVE_self_ or single PVE_nei_ mostly overrepresented the actual amounts of PVE_self_ or PVE_nei_, respectively (Fig. S3). The overrepresentation of the single PVE_self_ and single PVE_nei_ was observed when either the self or neighbor effects were absent in the simulation (Fig. S4). These results indicate that (i) single PVE_nei_ should not be used, (ii) partial PVE_nei_ suffered from its collinearity with the single PVE_self_, and (iii) net PVE_nei_ provides a conservative estimate for the genome-wide contribution of neighbor effects to phenotypic variation.

**Figure 3.**
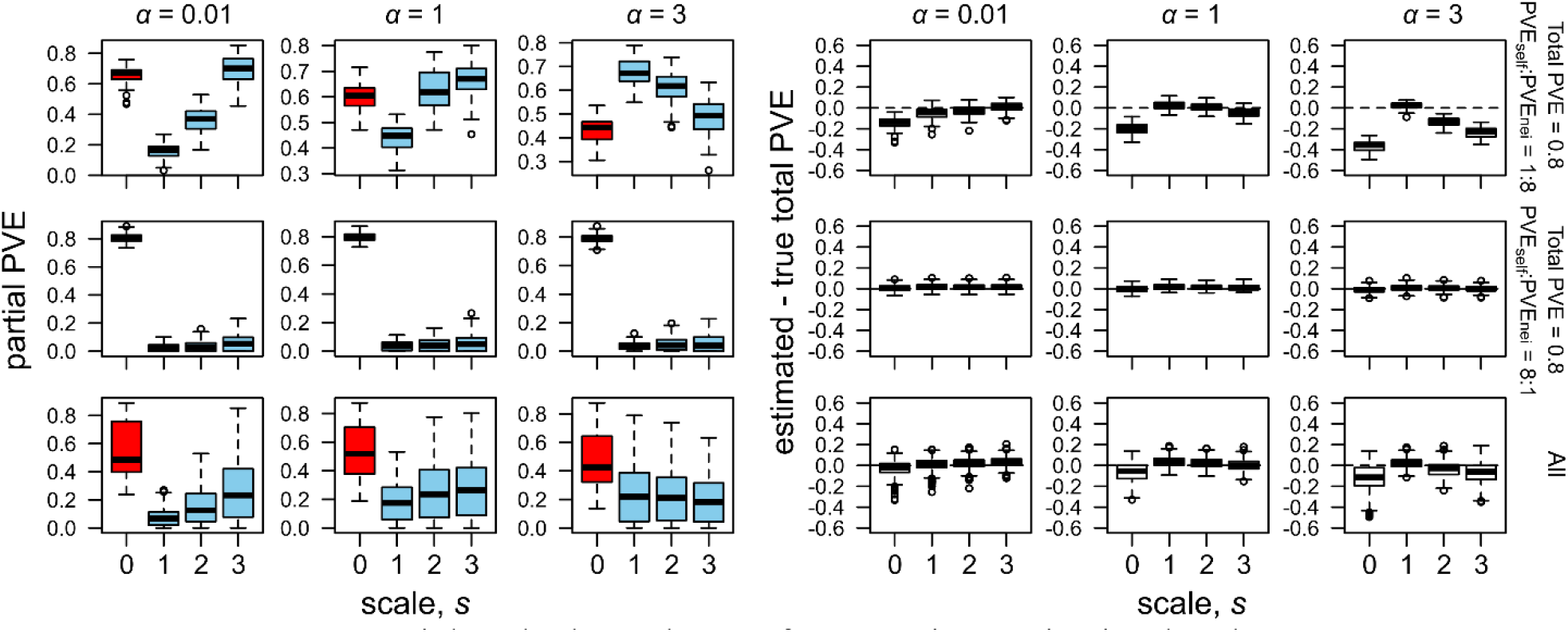
Spatial scale dependence of PVE estimates in simulated phenotypes. The broad, intermediate, and narrow effective range of neighbor effects are represented by weak (*α* = 0.01), moderate (*α* = 1), and strong (*α* = 3) distance decay coefficients, respectively. Partial PVE (left) and the accuracy of the total PVE estimation (right) are shown along the spatial scale from the first nearest (*s* = 1) to the third nearest (*s* = 3) neighbors, with distinct relative contributions of the self and neighbor effects to a phenotype (PVE_self_:PVE_nei_ = 1:8 or 8:1). Boxplots show center line: median, box limits: upper and lower quartiles, whiskers: 1.5 × interquartile range, and points: outliers. In the left panels, red boxes indicate partial PVE_self_ at *s* = 0 (corresponded to single PVE_self_), while blue boxes indicate partial PVE_nei_ at *s* ≠ 0. In the right panels, horizontal dashed lines indicate a perfect match between the estimated and true total PVE.

**Table 1.**
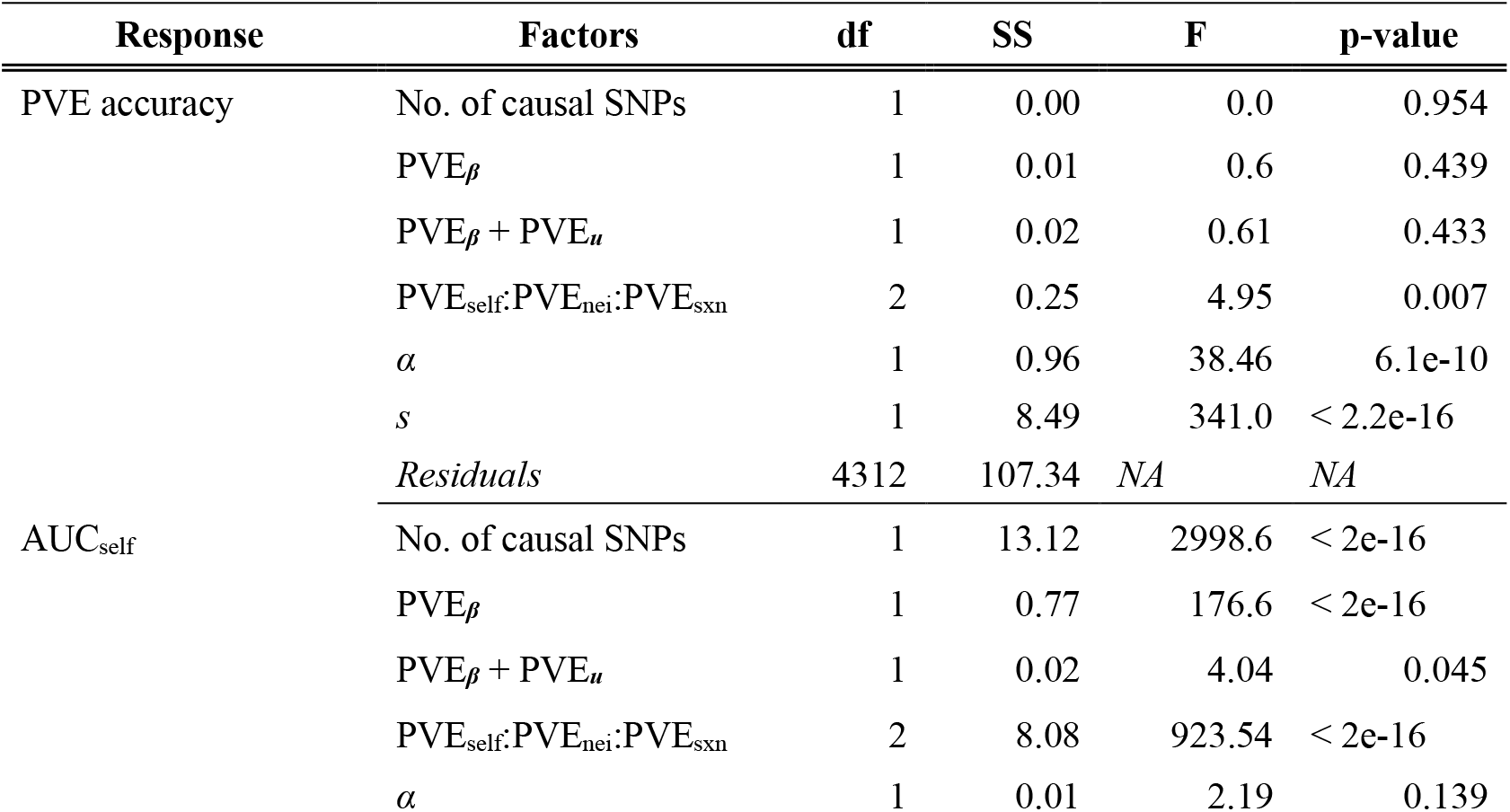
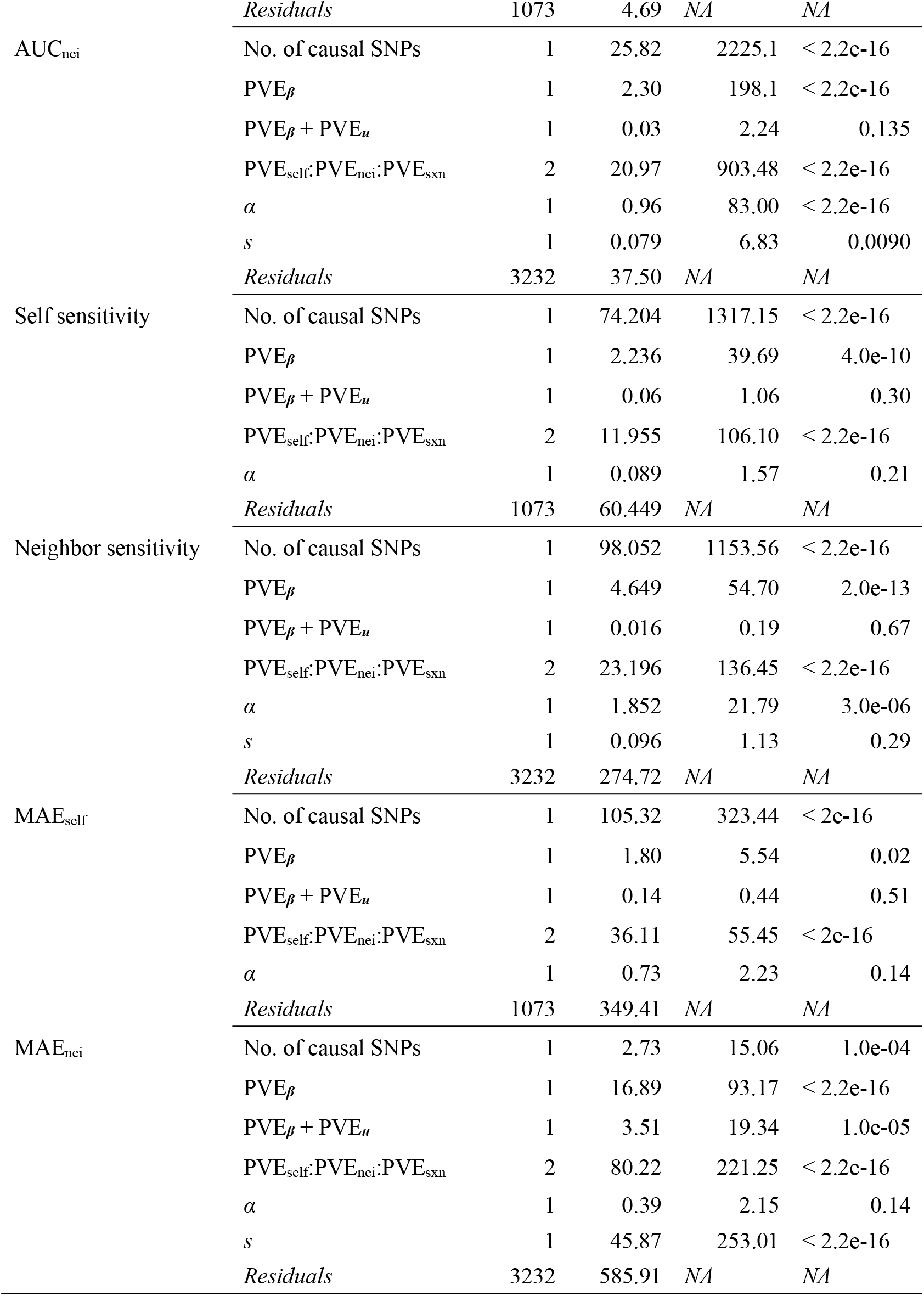
Factors affecting variance estimation and causal variant detection in the simulated phenotypes. The accuracy of the proportion of the phenotypic variation explained (PVE) was defined as the PVE accuracy = (estimated total PVE – true total PVE) / true total PVE. The power was represented by the area under the ROC curve (AUC). The sensitivity to detect self or neighbor effects was evaluated using the true positive rate of the ROC curve, when the false positive rate = 0.05. The accuracy of the effect size estimates were evaluated using the mean absolute errors (MAE) between the true and estimated fixed effects. ANOVA tables show the degree of freedom (df), sum of squares (SS), *F*-statistics, and *p*-values. Explanatory factors are the number of causal SNPs, proportion of phenotypic variation explained (PVE) by major-effect genes (PVE_***β***_), total PVE by major-effect genes and variance components (PVE_***β***_ + PVE_***u***_), relative contribution of self, symmetric, and asymmetric neighbor effects (PVE_self_:PVE_nei_:PVE_sxn_), and distance decay coefficient α. For the neighbor effects, the difference of the reference spatial scales (*s* = 1 - 3) was also considered an explanatory variable. NA means not available.

Although the partial PVE_nei_ could not be used to quantify the net contribution of the neighbor effects, this metric inferred spatial scales at which neighbor effects remained effective. If the distance decay was weak (small value of decay coefficient *α*) and the effective range of the neighbor effects was broad, partial PVE_nei_ increased linearly as the reference spatial scale was broadened (Fig. 3). On the other hand, if the distance decay was strong (large value of decay coefficient *α*) and the effective scale of the neighbor effects was narrow, partial PVE_nei_ decreased as the reference spatial scale was broadened or saturated at the scale of the first nearest neighbors (Fig. 3). Considering the spatial dependency of the partial PVE_nei_, we could estimate the effective spatial scales by *Δ*PVE_nei_ = partial PVE_nei,*s*+1_ – partial PVE_nei,*s*_ and by the scale that resulted in the maximum *Δ*PVE_nei_ as *s* = arg max *Δ*PVE_nei_ (Fig. S5).

The spatial scale that yielded the maximum AUC for neighbor effects, coincided with the patterns of the partial PVE_nei_ across the range of *s*. If the distance decay was weak (*α* = 0.01) and the effective range of the neighbor effects was broad, the AUC_nei_ increased linearly as the reference spatial scale was broadened (Fig. 4). If the distance decay was strong (large value of decay coefficient *α*) and the effective scale of the neighbor effects was narrow, the AUC_nei_ did not increase even when the reference spatial scale was broadened (Fig. 4). Thus, the first nearest scale was enough to detect neighbor signals, unless the distance decay was very weak.

**Figure 4.**
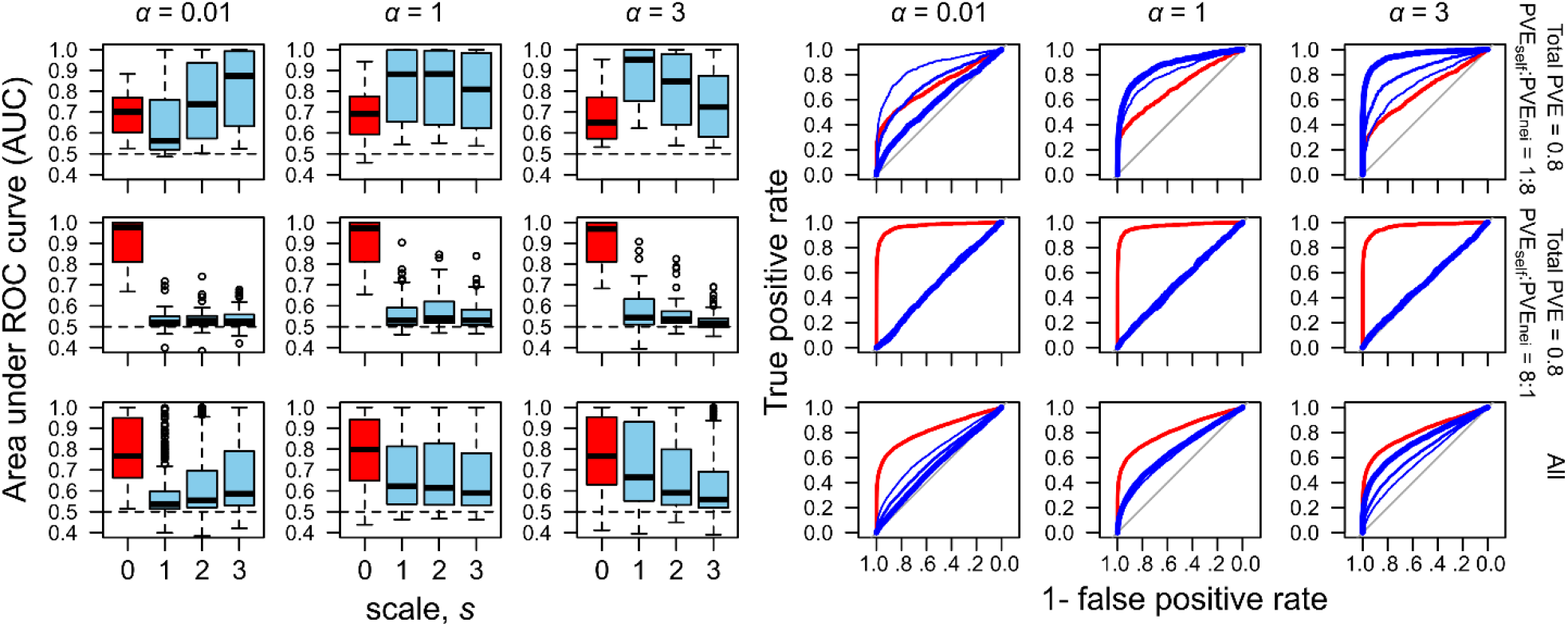
Spatial scale dependence of the power to detect causal SNPs in simulated phenotypes. The broad, intermediate, and narrow effective range of neighbor effects are represented by weak (*α* = 0.01), moderate (*α* = 1), and strong (*α* = 3) distance decay coefficients, respectively. Receiver operating characteristic (ROC) curves (right) and the area under the ROC curve (AUC) (left) are shown alongside the spatial scales from the first nearest (*s* = 1) to the third nearest (*s* = 3) neighbors, with the distinct relative contributions of the self and neighbor effects to a phenotype (PVE_self_ :PVE_nei_ = 1:8 or 8:1). Red boxes and curves indicate self-effects, while blue boxes indicate neighbor effects. The thickness of the blue curves indicates reference spatial scales as follows: *s* = 1 (thick), 2 (medium), or 3 (thin). The horizontal dashed lines in the left panels indicates that the AUC = 0.5, i.e., no detection of causal variants. The ROC curves in the right panels are depicted based on ten iterations with 50 causal SNPs.

In terms of the AUC, we also found that the number of causal SNPs, the amount of PVE by neighbor effects (controlled by the total PVE = PVE***β*** + PVE_***u***_; and ratio of PVE_self_:PVE_nei_), and the distance decay coefficient *α* were significant factors affecting the power to detect neighbor signals (Table 1). The power to detect self-genotype effects depended on the number of causal SNPs and PVE*β* but was not significantly influenced by the distance decay coefficient of the neighbor effects (Table 1). The power to detect self-genotype signals changed from strong (AUC_self_>0.9) to weak (AUC_self_<0.6), depending on the number of causal SNPs, the PVE by the major-effect genes, and as the relative contribution from the PVE_self_ increased (Fig. S6). Compared to the self-genotype effects, it was relatively difficult to detect neighbor effects (Fig. 4; Fig. S6), ranging from strong (AUC_nei_>0.9) to little (AUC_nei_ near to 0.5) power. When the number of causal SNPs = 10, the power to detect neighbor signals decreased from high (AUC_nei_>0.9) to moderate (AUC_nei_>0.7) with the decreasing PVE_***β***_ and the distance decay coefficient (Fig. 4; Fig. S6). There was almost no power to detect neighbor signals (AUC_nei_ near to 0.5) when the number of causal SNPs = 50 and PVE_nei_ had low contributions (Fig. S6). The result of the simulations indicated that strong neighbor effects were detectable when a target trait was governed by several major genes and the range of neighbor effects was spatially limited. Additionally, linear mixed models outperformed standard linear models as there were 8.8% and 1.4% increases in their power to detect self and neighbor signals, respectively (AUC_self_, paired *t*-test, mean of the difference = 0.088, *p*-value<10^-16^; AUC_nei_, mean of the difference = 0.014, *p*-value<10^-16^). When the neighbor phenotype 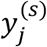. was incorporated instead of the genotype 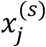, the power to detect neighbor effects was very weak, such that the AUC_nei_ decreased to almost 0.5.

To examine misclassifications between the self and neighbor signals, we compared the sensitivity, effect size estimates, and *p*-value scores among causal SNPs having non-zero coefficients of the true *β*_1_ and *β*_2_. The sensitivity to detect the self and neighbor effects was largely affected by the number of causal SNPs, the amount of PVE by the major-effect genes PVE_***β***_, and the relative contribution of the self and neighbor effects (controlled by PVE_self_:PVE_nei_) (Table 1; Fig. S7). The mean absolute errors of the self-effect estimates for 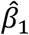 largely depended on the number of causal SNPs and the relative contribution of the variance components, while those of the neighbor effect estimates for 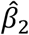 were dependent on the relative contribution of the variance components and the spatial scales to be referred (Table 1; Fig. S8). Given that the self and neighbor signals were sufficiently detected when the number of causal SNPs was 50 (Fig. S6), *p*-values under this condition were compared between the causal and non-causal SNPs. We observed that strong self-signals (*β*_1_ ≠ 0) were unlikely to be detected as neighbor effects (Fig. 5). Causal SNPs responsible for both the self and neighbor effects (*β*_1_ ≠ 0 and *β*_2_ ≠ 0) were better detected than the non-causal SNPs (*β*_1_ = 0 and *β*_2_ = 0). The sensitivity to detect neighbor effects was large when the true contribution of the neighbor effects was as large as PVE_self_ :PVE_nei_ = 1:8, but decreased when the contribution of the self-effects was as large as PVE_self_ :PVE_nei_ = 8:1 (Fig. S7). In contrast, if the contribution of the neighbor effects was relatively large (PVE_self_ :PVE_nei_ = 1:8), the SNPs responsible for the neighbor effects alone (*β*_1_ = 0 and *β*_2_ ≠ 0), could also be detected as self-effects (Fig. 5). As expected by the level of collinearity (Fig. S2), the false positive detection of the self -effects was more likely when the distance decay coefficient was small, and the effective range of the neighbor effects was broad (Fig. 5). This coincided with the strength of the collinearity (Fig. S2), as the false positive detection of self-effects and false negative detection of neighbor effects are more likely if the MAF was small (Fig. S9). Consistent with the false positive detection, the sensitivity to detect self -effects remained large, even when the contribution of the neighbor effects was far larger (Fig. S7). Strong self-effects (*p*-value < 10^-5^ for 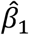) and slight neighbor effects (*p*-value < 0.05 for 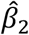 at *s* = 1 and *α* = 3) remained when asymmetric neighbor effects were strong (*β*_1_ ≠ 0 and *β*_2_ ≠ 0 and *β*_12_ ≠ 0 and PVE_self_:PVE_nei_:PVE_sxn_ = 1:1:8; Fig. S10). These results indicate that (i) the collinearity may lead to the false positive detection of self-effects, yet is unlikely to result in the false positive detection of neighbor effects, and that (ii) smaller MAFs are more likely to cause the false positive detection of self-effects and decrease the power to detect true neighbor effects.

**Figure 5.**
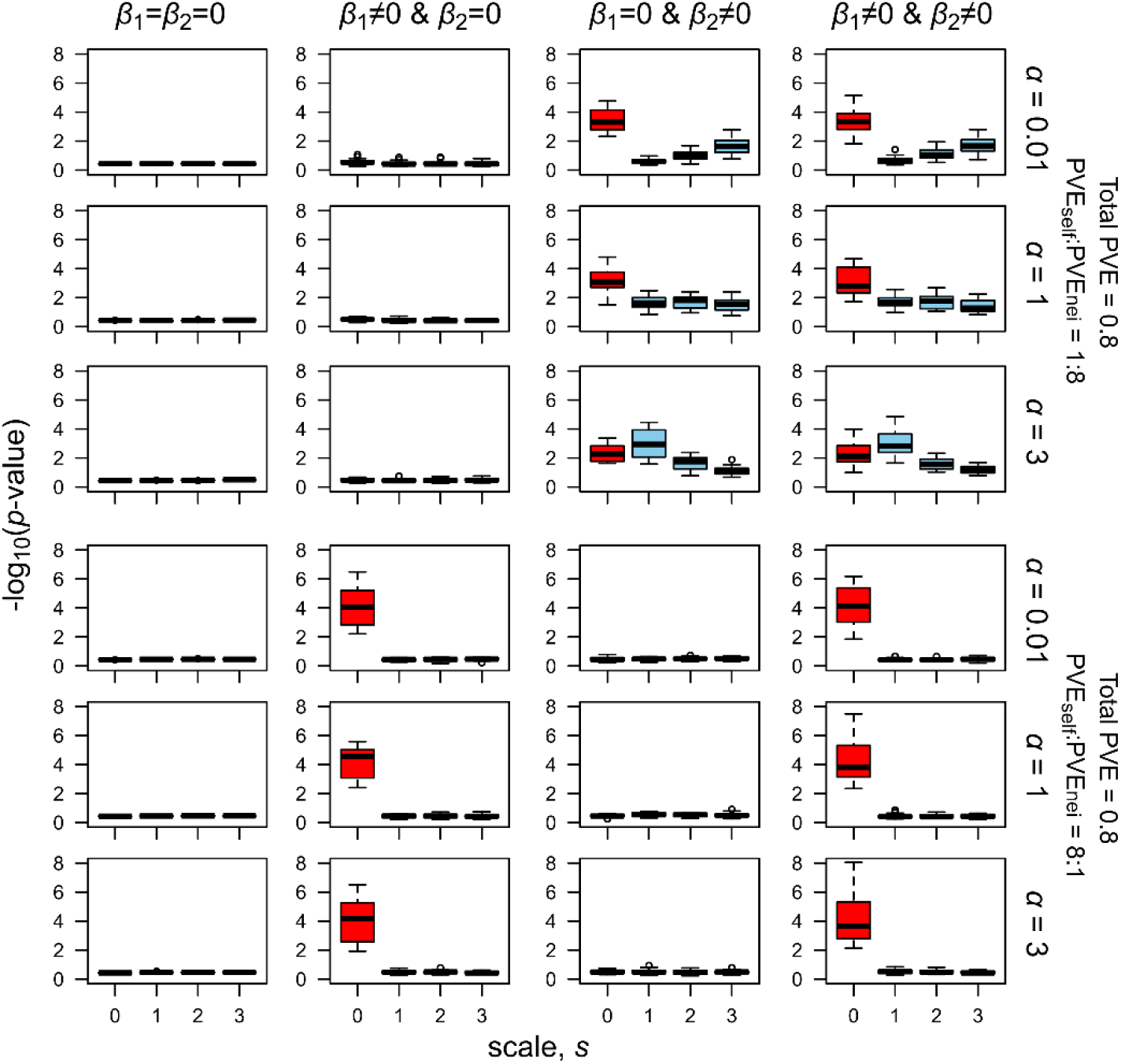
Signals of the self and neighbor effects when either the self or neighbor effects were for 50 causal SNPs. The score of −log_10_(*p*-value) is averaged within each iteration and is shown for the non-causal SNPs (*β*_1_ = *β*_2_ = 0), SNPs responsible for self-effects alone (*β*_1_ ≠ 0 and *β*_2_ = 0), SNPs responsible for neighbor effects alone (*β*_1_ = 0 and *β*_2_ ≠ 0), and SNPs responsible for both self and neighbor effects (*β*_1_ ≠ 0 and *β*_2_ ≠ 0). Red and blue boxes show −log_10_(*p*-value) distributions among the iterations for the self and neighbor effects, respectively.

### *Arabidopsis* herbivory data

To estimate PVE_self_ and PVE_nei_, we applied a linear mixed model (eq. 3) to the leaf damage score data for the field-grown *A. thaliana.* The leaf damage variation was significantly explained by the single PVE_self_ that represented additive genetic variation (single PVE_self_ = 0.173, 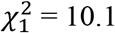, *p*-value = 0.005). Variation partitioning showed a significant contribution of neighbor effects to the phenotypic variation in the leaf damage at the nearest scale (partial PVE_nei_ = 0.214, 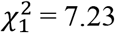, *p*-value = 0.004 at *s* = 1: Fig. S11). The proportion of phenotypic variation explained by the neighbor effects did not increase when the neighbor scale was referred up to the nearest and second nearest individuals (partial PVE_nei_ = 0.14, 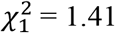, *p*-value = 0.166 at *s* = 2: Fig. S11); therefore, the effective scale of the neighbor effects was estimated at *s* = 1 and variation partitioning was stopped at *s* = 2. These results indicated that the effective scale of the neighbor effects on the leaf damage was narrow (*s* = 1) and the net PVE_nei_ at *s* = 1 explained an additional 8% of the PVE compared to the additive genetic variation attributable to the single PVE_self_ (Fig. 6a). The genotype data had moderate to strong element-wise correlation between ***K***_1_ and ***K***_2_ in these analyses (*r* = 0.60 and 0.78 at *s* = 1 and 2 among 199 accessions with eight replicates). We additionally incorporated the neighbor phenotype 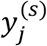 instead of the neighbor genotype 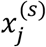 in eq. 2, but the partial PVE_nei_ did not increase (partial PVE_nei_ = 0.066 and 0.068 at *s* = 1 and 2, respectively).

**Figure 6.**
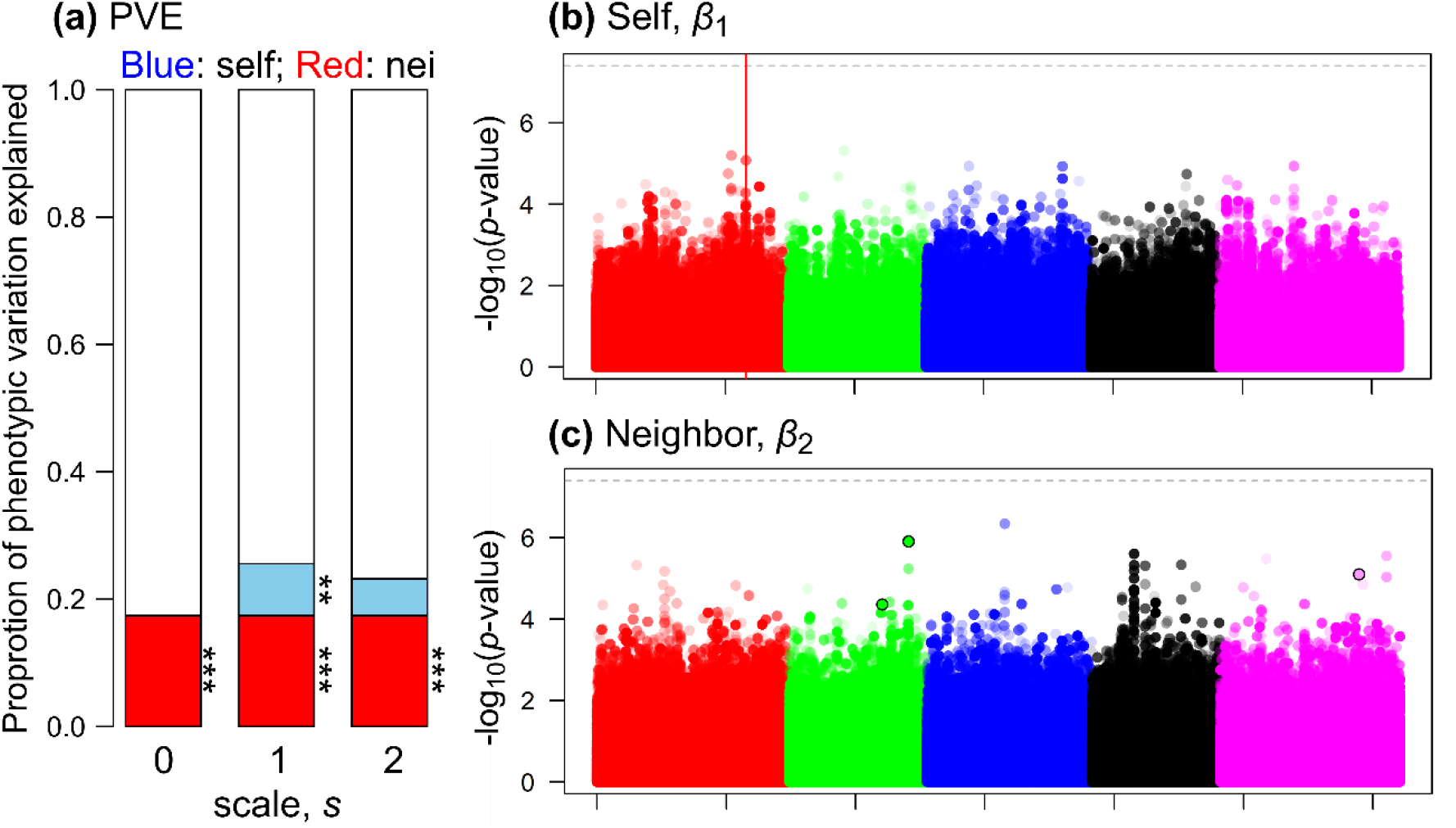
Pilot GWAS of leaf damage scores on field-grown *Arabidopsis thaliana.* (a) Proportion of phenotypic variation explained (PVE) by the self-genotype (red) or neighbor effects (blue). The PVE_self_ was represented by the single PVE_self_ that represented additive genetic variance, while the net contribution of the neighbor effects was evaluated using the net PVE_nei_ = total PVE – single PVE_self_. Asterisks highlight a significant fraction with stepwise likelihood ratio tests, from simpler to complex models: \*\**p*-value < 0.01: \*\*\**p*-value < 0.001. (b, c) Manhattan plots for the self or neighbor effects. The first to fifth chromosomes are differently colored, where lighter plots indicate smaller MAF. Horizontal dashed lines indicate the threshold after Bonferroni correction at *p*-value < 0.05. The red vertical line in panel (a) indicates an SNP position near the *GS-OX2* locus, while the three circles highlighted by a black outline in panel (b) indicates the variants subject to the post hoc simulation (Fig. 7). Results of the self and neighbor effects are shown at *s* = 0 (i.e., standard GWAS) and *s* = 1, respectively.

The standard GWAS of the self-genotype effects on the leaf damage detected the SNPs with the second and third largest −log_10_(*p*-values) scores, on the first chromosome (chr1), though they were not above the threshold for Bonferroni correction (Fig. 6b; Table S2). The second SNP at chr1-21694386 was located within ~10 kb of the three loci encoding a disease resistance protein (CC-NBS-LRR class) family. The third SNP at chr1-23149476 was located within ~10 kb of the AT1G62540 locus that encodes a flavin-monooxygenase glucosinolate S-oxygenase 2 (FMO GS-OX2). No GOs were significantly enriched for the self-effects on herbivory (false discovery rate > 0.08). A QQ-plot did not exhibit an inflation of *p*-values for the self-genotype effects (Fig. S12).

Regarding the neighbor effects on leaf damage, we found non-significant but weak peaks on the second and third chromosomes (Fig. 6c; Table S2). The second chromosomal region had higher association scores than those predicted by the QQ-plot (Fig. S12). A locus encoding FAD-binding Berberine family protein (AT2G34810 named *BBE16*), which is known to be up-regulated by methyl jasmonate (Devoto et al. 2005), was located within the ~10 kb window near SNPs with the top eleven −log_10_(*p*-values) scores on the second chromosome. Three transposable elements and a pseudogene of lysyl-tRNA synthetase 1 were located near the most significant SNP on the third chromosome. No GOs were significantly enriched for the self-effects on herbivory (false discovery rate > 0.9). We additionally tested the asymmetric neighbor effects of *β*_12_ in the real dataset on field herbivory, but the top 0.1% of the SNPs for the neighbor effects for *β*_2_, did not overlap with those of the asymmetric neighbor effects *β*_12_ (Table S2).

Based on the estimated coefficients 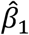 and 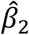, we ran a post hoc simulation to infer a spatial arrangement that minimizes a population sum of the leaf damage ∑*y_i_* = *β*_1_∑*x_i_* + *β*_2_ ∑_<*i,j*>_ *x_i_x_j_*. The constant intercept *β*_0_, the variance component *u_i_*, and residual *e_i_* were not considered because they were not involved in the deterministic dynamics of the model. Figure 7 shows three representatives and a neutral expectation. For example, a mixture of a dimorphism was expected to decrease the total leaf damage for an SNP at chr2-14679190 near the *BBE16* locus (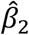: Fig. 7a). On the other hand, a clustered distribution of a dimorphism was expected to decrease the total damage for an SNP at chr2-9422409 near the AT2G22170 locus encoding a lipase/lipooxygenase PLAT/LH2 family protein (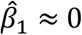, 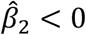: Fig. 7b). Furthermore, a near monomorphism was expected to decrease the leaf damage for an SNP at chr5-19121831 near the AT5G47075 and AT5G47077 loci encoding low-molecular cysteine-rich proteins, LCR20 and LCR6 (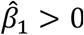,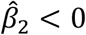: Fig. 7c). If the self and neighbor coefficients had no effects, we would observe a random distribution and no mitigation of damage i.e., ∑*y_i_* ≈ 0 (Fig. 7d). These post hoc simulations suggested a potential application for neighbor GWAS for the optimization of the spatial arrangements in field cultivation.

**Figure 7.**
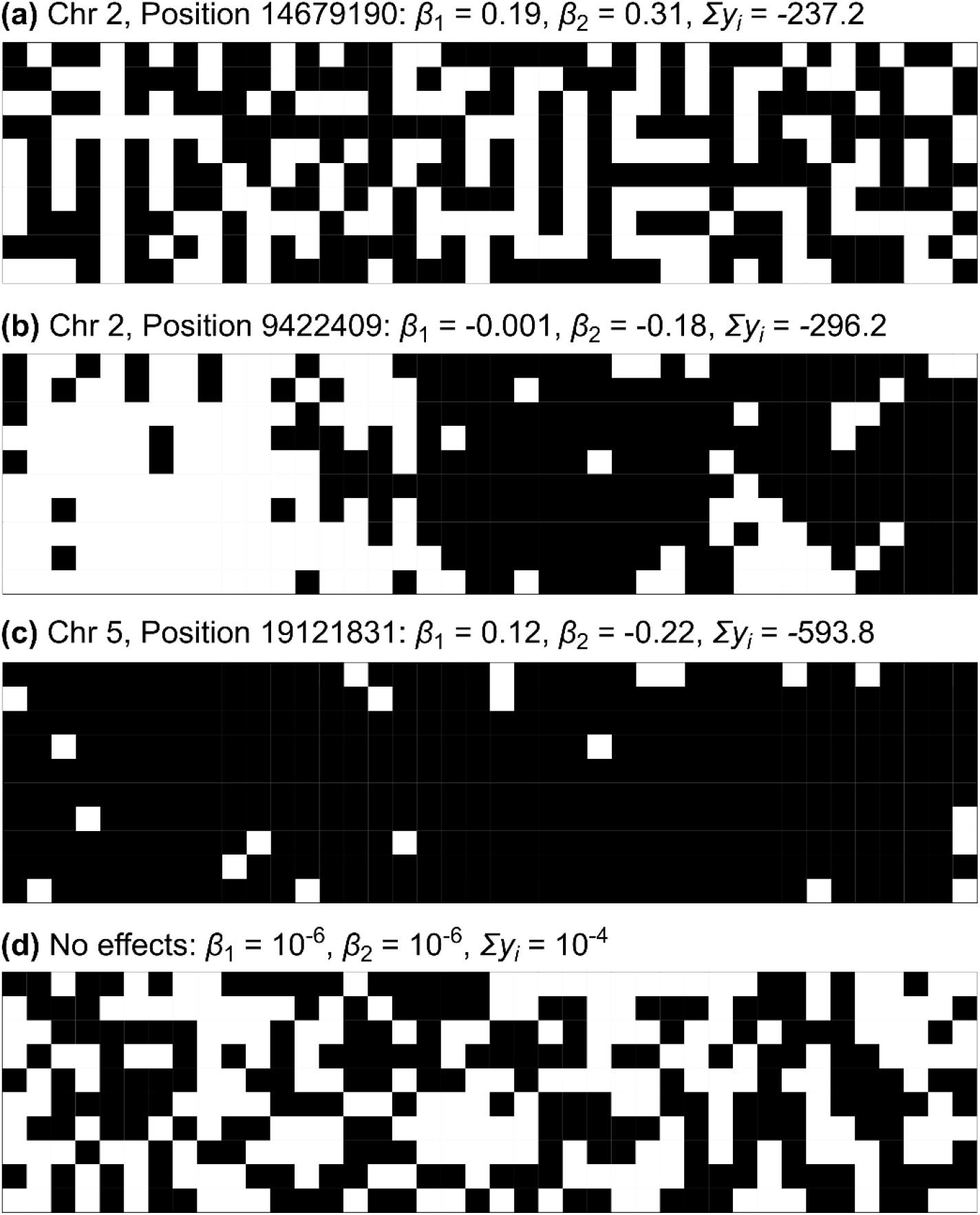
Post hoc simulations exemplifying a spatial arrangement of the two alleles expected by the estimated self and neighbor effects, 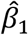 and 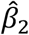, on the leaf damage score of *Arabidopsis thaliana.* The population sum of the leaf damage ∑*y_i_*, = *β*_1_∑*x_i_* + *β*_2_ ∑_<*i,j*>_*x_i_x_j_* was minimized using 1000 iterations of simulated annealing from a random distribution of two alleles in a 10 × 40 space.

### Comparing self *p*-values between the neighbor GWAS and GEMMA

To ascertain whether the self-genotype effects in the neighbor GWAS agree with those of a standard GWAS, we compared the *p*-value scores between the rNeighborGWAS package and the commonly used GEMMA program (Fig. S13). For the leaf damage score, the neighbor GWAS yielded almost the same −log_10_(*p*-values) scores for the self-effects as the GEMMA program (*r* = 0.9999 among all the 1,242,128 SNPs). The standard GWAS, using the flowering time phenotype, also yielded the consistent −log_10_(*p*-values) scores between the neighbor GWAS and GEMMA (*r* = 0.9999 among all the 1,814,755 SNPs: Fig. S13). Both the flowering time GWAS using the neighbor GWAS and GEMMA found two significant SNPs above the genome-wide Bonferroni threshold on chromosome 5 (chr5-18590741 and chr5-18590743, MAF = 0.49 and 0.49, −log_10_(*p*-value) = 7.797 and 7.797 for the neighbor GWAS; chr5-18590741 and chr5-18590743, MAF = 0.49 and 0.49, −log_10_(*p*-value) = 7.798 and 7.798 for GEMMA), which were located within the *Delay of Germination 1* (*DOG1*) locus, that was reported previously by Alonso-Blanco et al. (2016). Another significant SNP was observed at the top of chromosome 4 (chr4-317979, MAF = 0.12, −log_10_(*p*-value) = 7.787 and 7.933 for the neighbor GWAS and GEMMA), which was previously identified as a quantitative trait loci underlying flowering time in long-day conditions (Aranzana et al. 2005).

## DISCUSSION

### Spatial and genetic factors underlying simulated phenotypes

Benchmark tests using simulated phenotypes revealed that appropriate spatial scales could be estimated using the partial PVE_nei_ of the observed phenotypes. When the scale of the neighbor effects was narrow or moderate (*α* = 1.0 or 3.0), the scale of the first nearest neighbors would be optimum for increasing the AUC to detect neighbor signals. In terms of the neighbor effects in the context of plant defense, mobile animals (e.g., mammalian browsers and flying insects) often select a cluster of plant individuals (e.g., Bergvall et al. 2006; Hambäck et al. 2009; Sato and Kudoh 2015; Verschut et al. 2016). In this case, the neighbor effects could not be observed among individual plants within a cluster (Sato and Kudoh 2015). The exponential distance decay at *α* = 0.01 represented situations in which the effective range of the neighbor effects was too broad to be detected; only in such situations should more than the nearest neighbors be referred to, to gain the power to detect neighbor effects. We also considered the asymmetric neighbor effects where the neighbor genotype similarity had significant effects on one genotype, but not on another genotype. In this situation, strong self-effects could be observed when the symmetric neighbor effects were weakened. This additional result suggests that asymmetric neighbor effects should be tested if strong self-effects and weak symmetric neighbor effects are both detected at a single locus.

Neighbor effects are more likely to contribute to phenotypic variation when its effective range becomes narrow due to a strong distance decay (*α* = 3), as suggested by the net PVE_nei_. However, the total phenotypic variation was explained relatively well by the single PVE_self_ that represented additive polygenic effects. Previous studies showed that genetic interactions could lead to an overrepresentation of narrow-sense heritability in GWAS (e.g., Zuk et al. 2012; Young and Durbin 2014). This occurs because the SNP heritability is represented by the genetic similarity between individuals, and thereby covariance of the kinship matrix helps to fit the phenotypic variance attributable to gene-by-gene interactions (Young and Durbin 2014; Schrauf et al. 2020). This problem is also observed in the neighbor GWAS that models pairwise interactions at a focal locus among neighboring individuals. Given the difficulty in distinguishing the kinship and genetic interactions, we conclude that the non-independence of the self and neighbor effects is an intrinsic feature of the neighbor GWAS, and that the difference of the PVE between a standard and neighbor GWAS i.e., net PVE_nei_ should be used as a conservative estimate of PVE_nei_.

### Neighbor GWAS of the field herbivory on *Arabidopsis*

Our genetic analysis of the neighbor effects is of ecological interest, as the question of how host plant genotypes shape variations in plant–herbivore interactions, is a long-standing question in population ecology (e.g., Karban 1992; Underwood and Rausher 2000; Utsumi et al. 2011). Despite the low PVE and several confounding factors under field conditions, the present study illustrated the significant contribution of neighbor genotypic identity, to the spatial variation of the herbivory on *A. thaliana*. Although the additional fraction explained by the neighbor effects was 8%, this amount was plausible in the GWAS of complex traits. For example, the variance components of epistasis explained 10-14% PVE on average for 46 traits in yeast (Young and Durbin 2014). Even when heritability is high, the significant variants have often explained a small fraction of PVE, which is known as the missing heritability problem in plants and animals (Brachi et al. 2011; López-Cortegano and Caballero 2019).

Regarding the self-genotype effects, we detected *GS-OX2* near the third top-scoring SNP on the first chromosome. GS-OX2 catalyzes the conversion of methylthioalkyl to methylsulfinylalkyl glucosinolates (Li et al. 2008) and is up-regulated in response to feeding by the larvae of the large white butterfly (*Pieris brassicae*) (Geiselhardt et al. 2013). On the other hand, the second top-scoring SNP of the neighbor effects was located near the *BBE16* locus, responsive to methyl jasmonate, a volatile organic chemical that is emitted from damaged tissue and elicits the defense responses of other plants (Reymond and Farmer 1998; van Poecke 2007). However, because none of the associations were significant above a genome-wide Bonferroni threshold, they should be interpreted cautiously. Nearby genes should only be considered candidates, and further work is necessary to confirm that they exert any neighbor effects on herbivory.

### Potential limitation

Despite many improvements, it is more difficult for GWAS to capture rare causal variants than common ones (Lee et al. 2014; Auer and Littre 2015; Bomba et al. 2018). This problem is more severe in neighbor GWAS, because smaller MAFs result in stronger collinearity between the self-genotype effects *x_i_* and the neighbor genotypic identity 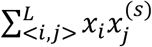. Our simulations showed that the rare variants responsible for the neighbor effects might be misclassified as self-effects, though the opposite was not found, i.e., the misclassification of self-signals into neighbor effects could be suppressed. In GWAS, genotype data usually contains minor alleles and possess kinship structures to some extent, making collinearity unavoidable. To anticipate false positive detection of neighbor effects, the significance of variance components and marker effects involving neighbor effects should always be compared using the standard GWAS model.

The present neighbor GWAS focused on single-locus effects and did not incorporate locus-by-locus interactions. Although it is challenging to integrate all the association tests for epistasis into GWAS (Gondro et al. 2013; Young and Durbin 2014), it is possible that multiple combinations among different variants govern neighbor effects. For example, neighbor effects on insect herbivory may occur due to the joint action of volatile-mediated signaling and the accumulation of secondary metabolites (Dicke and Baldwin 2010; Erb 2018). The linear mixed model could be extended as exemplified by the asymmetric neighbor effects; however, we need to reconcile multiple criteria including the collinearity of explanatory variables, inflation of *p*-values, and computational costs. Further customization is warranted when analyzing more complex forms of neighbor effects.

## Conclusion

Based on the newly proposed methodology, we suggest that neighbor effects are an overlooked source of phenotypic variation in field-grown plants. GWAS have often been applied to crop plants (Jannink et al. 2010; Hamblin et al. 2011), where genotypes are known, and individuals are evenly transplanted in space. Considering this outlook for agriculture, we provided an example of neighbor GWAS across a lattice space in this study. However, wild plant populations sometimes exhibit more complex spatial patterns than those expected by the Ising model (e.g., Kizaki and Katori 1999; Schlicht and Iwasa 2004). In the rNeighborGWAS package, we allowed neighbor GWAS for a continuous two-dimensional space. While its application has now been limited to experimental populations, neighbor GWAS has the potential for compatibility with the emerging discipline of landscape genomics (Bragg et al. 2015). In this context, the additional R package could help future studies to test self and neighbor effects using a wide variety of plant species.

Neighbor GWAS may also have the potential to help determine optimal spatial arrangements for plant cultivation, as suggested by the post hoc simulation. Genome-wide polymorphism data are useful not only for identifying causal variants in GWAS, but also for predicting the breeding values of crop plants for genomic selection (e.g., Jannink et al. 2010; Hamblin et al. 2011; Yamamoto et al. 2017). Given that the neighbor GWAS consists of a marker-based regression, this methodology could also be expanded as a genomic selection tool to help predict population-level phenotypes in spatially structured environments.

## ACKNOWLEDGEMENTS

The authors would like to thank Ü. Seren, A. Korte, and M. Nordborg for kindly providing the full imputed SNP data prior to it being publicly available; T. Tsuchimatsu, K. Iwayama, and J. Bascompte for discussions; and Dynacom Co., Ltd. for technical assistance with the R package development. This study was supported by the Japan Science and Technology Agency (JST) PRESTO (Grant number, JPMJPR17Q4 and JPMJPR16Q9) to Y.S. and E.Y; Japan Society for the Promotion of Science (JSPS) Postdoctoral Fellowship (16J30005) to Y.S.; MEXT KAKENHI (18H04785) and the Swiss National Science Foundation to K.K.S.; URPP Global Change and Biodiversity of the University of Zurich to YS. and K.S.S.; and JST CREST (JPMJCR15O2 and JPMJCR16O3) to A.J.N. and K.K.S. The field experiment was supported by the Joint Usage/Research Grant of Center for Ecological Research, Kyoto University, Japan.

## DATA ARCHIVING

The leaf damage data on *A. thaliana* are included in the supporting information (Table S1). The simulation code and R script used in this study are available at the GitHub repository (https://github.com/naganolab/NeighborGWAS). R package version of the neighbor GWAS method is available at CRAN (https://cran.r-project.org/package=rNeighborGWAS).

