## Supplementary Figures for "Neighbor GWAS: incorporating neighbor genotypic identity into genome-wide association studies of field herbivory"

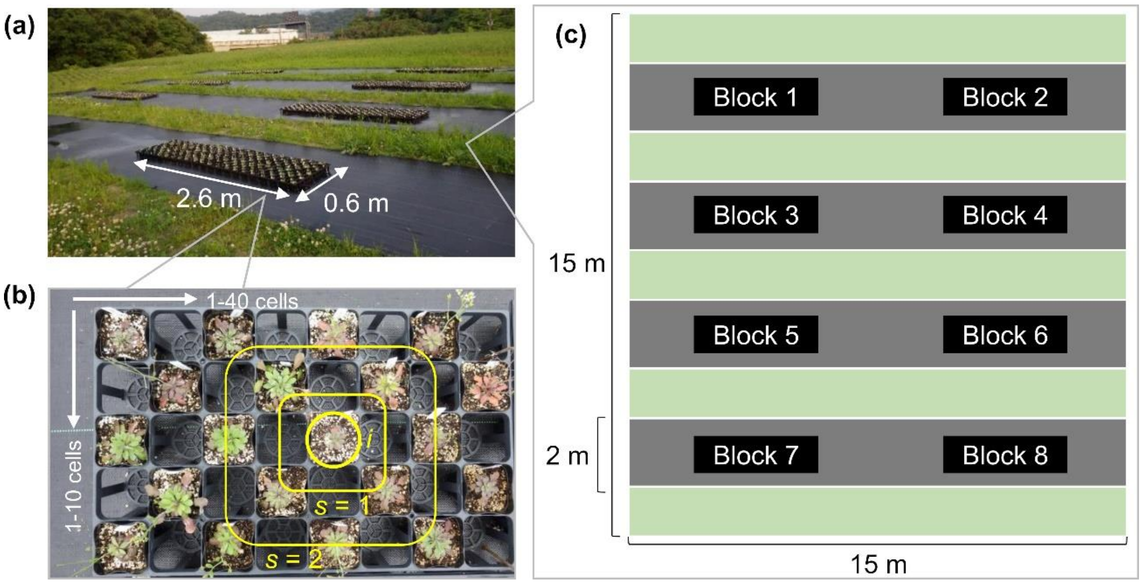


**Figure S1.** Experimental setting for the *Arabidopsis* herbivory data. (a) Photograph of the field site. Each 0.6 × 2.6 m block included a replicate of 199 accessions and an additional Col-0. (b) *Arabidopsis thaliana* plants were arranged in a 10 × 40 cell plastic tray. The plants were placed in a checkered pattern so that alternate cells were left empty; therefore, each of the 40 rows contained 5 plants and each of the 10 columns contained 20 plants. Yellow lines represent the *s*-th neighbor scales from a focal *i*-th plant. (c) A graphical explanation of the experimental area. A meadow (green) was separately covered with weed-masking sheets (grey).

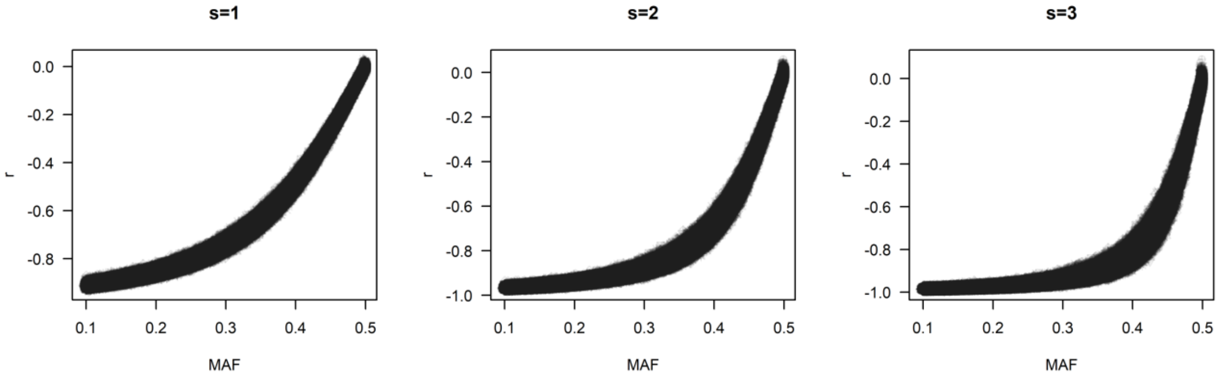


**Figure S2.** Dependency of the collinearity on the minor allele frequency (MAF) and spatial scales. Pearson correlation coefficients between the self-genotypic variable $x_i$ and the neighbor variables $(\sum_{<i,j>}^{L} x_i x_j^{(s)})/L$ are plotted against the MAF. The left, middle, and right panel show the cases for s = 1, 2, and 3, respectively. Shown are the results of ten iterations for random spatial distributions across a 36 × 36 lattice space, occupied by 1,206 *Arabidopsis thaliana* accessions. Partial genotypes were compiled from the RegMap panel (see the Method section in the main text).

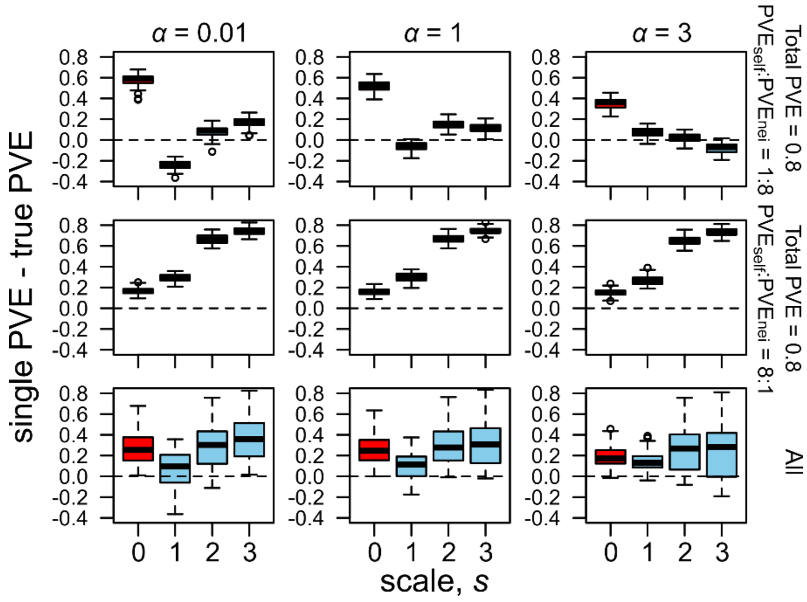


**Figure S3.** Accuracy of the single $PVE_{self}$ (red) and single $PVE_{nei}$ (blue) across different spatial scales in simulated phenotypes. The broad, intermediate, and narrow effective ranges of the neighbor effects are represented by weak ($\alpha$ = 0.01), moderate ($\alpha$ = 1), and strong ($\alpha$ = 3) distance decay coefficients, respectively. Discrepancies between the single PVE and true PVE are shown along the spatial scale from the first nearest ($s$ = 1) to the third nearest ($s$ = 3) neighbors, with distinct relative contributions of the self and neighbor effects to a phenotype ($PVE_{self}$:$PVE_{nei}$ = 1:8 or 8:1). Horizontal dashed lines indicate a perfect match between the estimated and true PVE by self or neighbor effects.

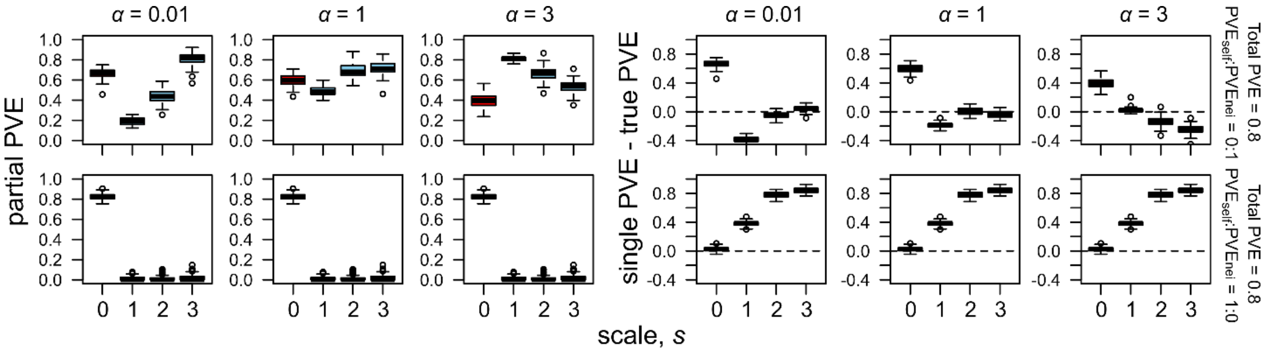


**Figure S4.** Partial and single PVE estimates when either self (lower panels) or neighbor (upper panels) effects determine a phenotype. The left panels show partial PVE estimates, the same as Figure 3, while the right panels show the accuracy of the single PVE estimates the same as Figure S3.

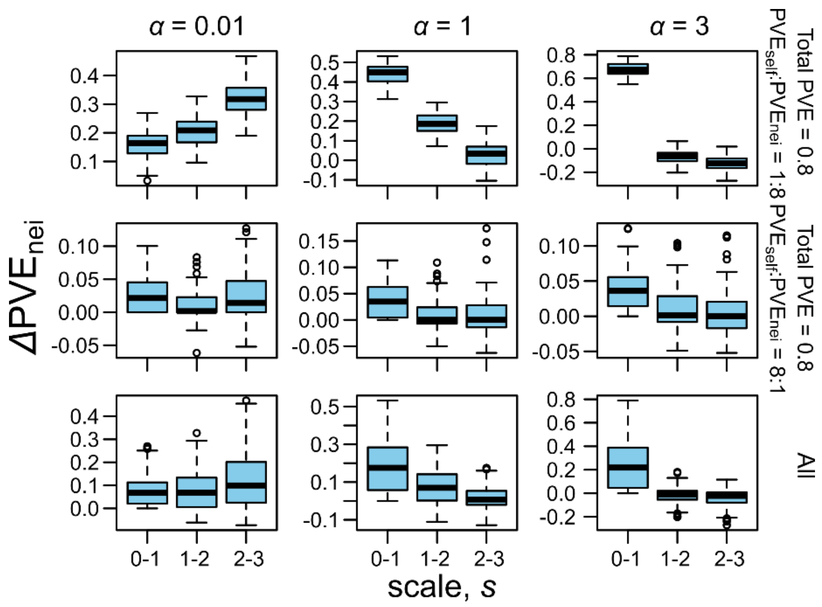


**Figure S5.** $\Delta$PVE metrics representing the shift of the partial PVE$_{nei}$ from the scale $s$ to $s$+1. The broad, intermediate, and narrow effective range of the neighbor effects are represented by weak ($\alpha$ = 0.01), moderate ($\alpha$ = 1), and strong ($\alpha$ = 3) distance decay coefficients, respectively. $\Delta$PVE$_{nei}$ for the s from 0 to 1 was equivalent with the partial PVE at $s$ = 1. $\Delta$PVE$_{nei}$ was largest at $s$ = 1, when the effective range of the neighbor effects was narrow or moderate. On the other hand, $\Delta$PVE$_{nei}$ was the largest at $s$ = 3 when the effective range of the neighbor effects was broad.

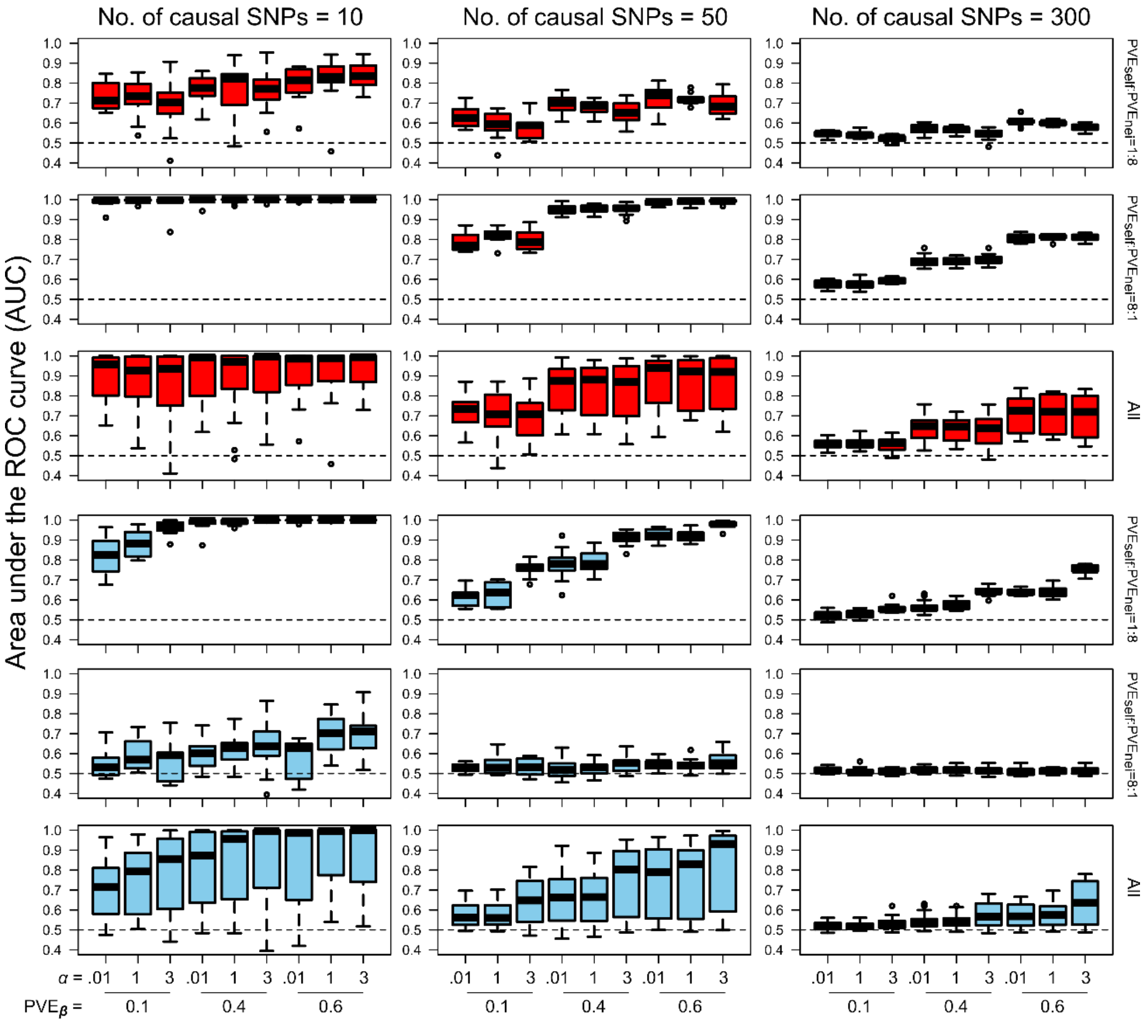


**Figure S6.** The power to detect the self and neighbor effects on the simulated phenotypes when the number of causal SNPs was 10, 50, or 300. The area under the ROC curve (AUC) is plotted against the distance decay coefficient $\alpha$ and the proportion of phenotypic variation explained by major-effect genes PVE$_{\beta}$ under the different relative contributions of self and neighbor effects. The AUC of the neighbor effects are shown at $s = 1$ for $\alpha = 3$; $s = 2$ for $\alpha = 1$; $s = 3$ for $\alpha = 0.01$, because these scales are known to give the maximum AUC among the three spatial scales (Fig. 3 and 4).

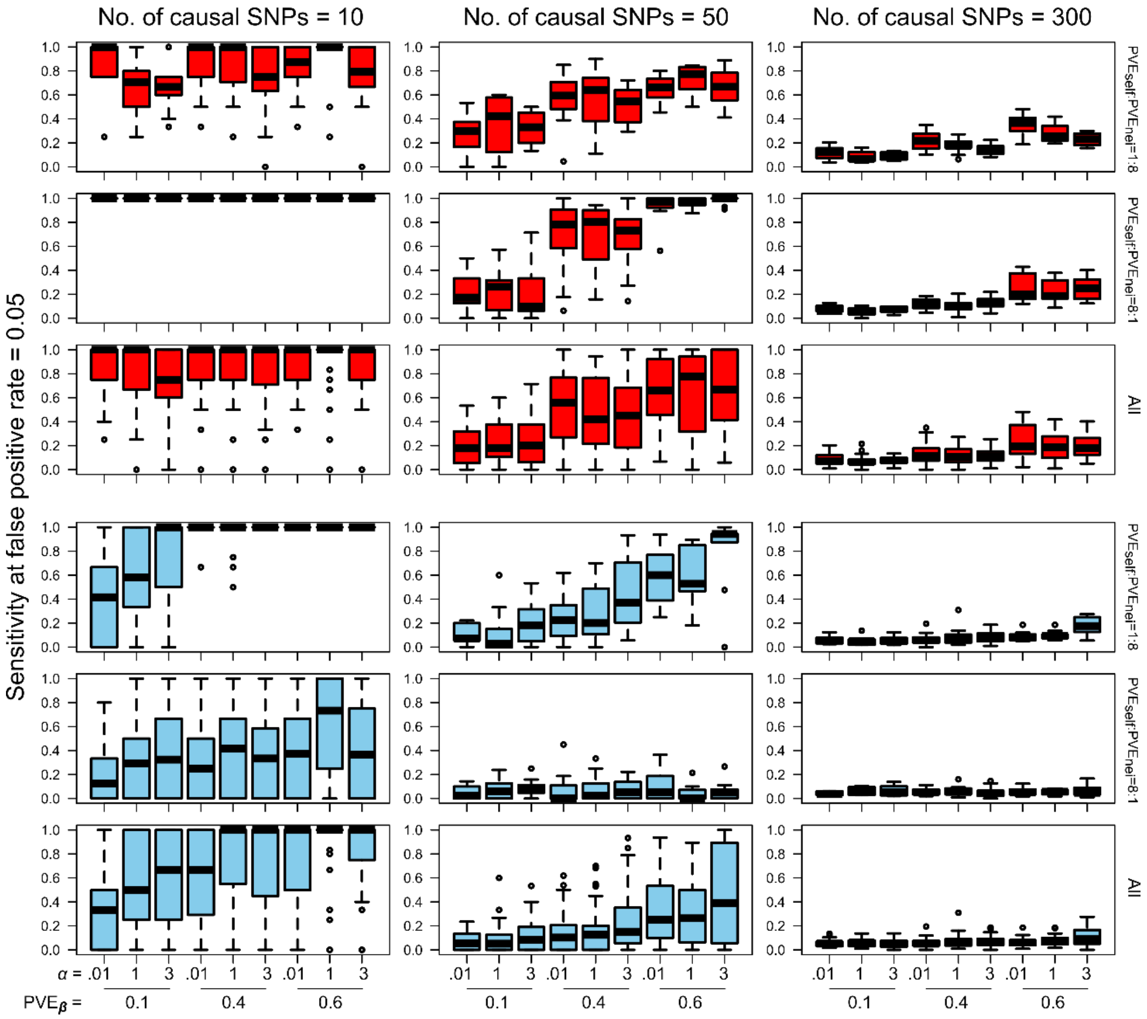


**Figure S7.** Sensitivity to detect self or neighbor signals when the number of causal SNPs was 10, 50, or 300. The sensitivity evaluated by the true positive rate when the false positive rate = 0.05, is plotted against the distance decay coefficient $\alpha$ and the proportion of phenotypic variation explained by the major-effect genes $\text{PVE}_{\beta}$ under different relative contributions of the self and neighbor effects. The sensitivity of neighbor effects are shown at $s = 1$ for $\alpha = 3$; $s = 2$ for $\alpha = 1$; and $s = 3$ for $\alpha = 0.01$, because these scales are known to give the maximum AUC among the three spatial scales (Fig. 3 and 4).

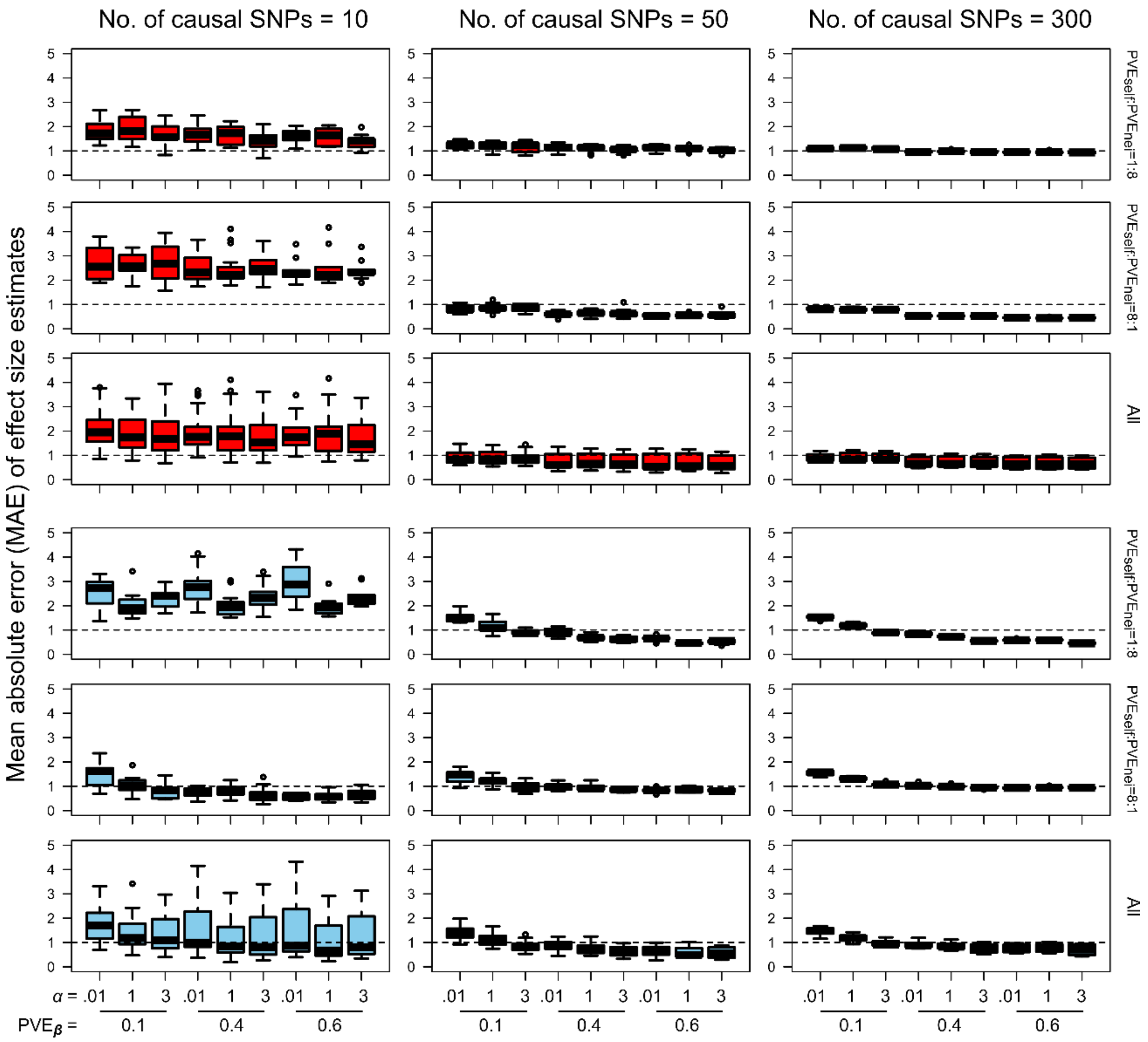


**Figure S8.** Mean absolute errors (MAE) between true and estimated $\beta$ when the number of causal SNPs was 10, 50, or 300. The MAE of the self-effect estimates $\hat{\beta}_1$ or neighbor effect estimates $\hat{\beta}_2$ is plotted against the distance decay coefficient $\alpha$ and the proportion of phenotypic variation explained by the major-effect genes $PVE_{\beta}$ under the different relative contributions of self and neighbor effects. MAE larger than 1.0 indicate larger errors than signals.

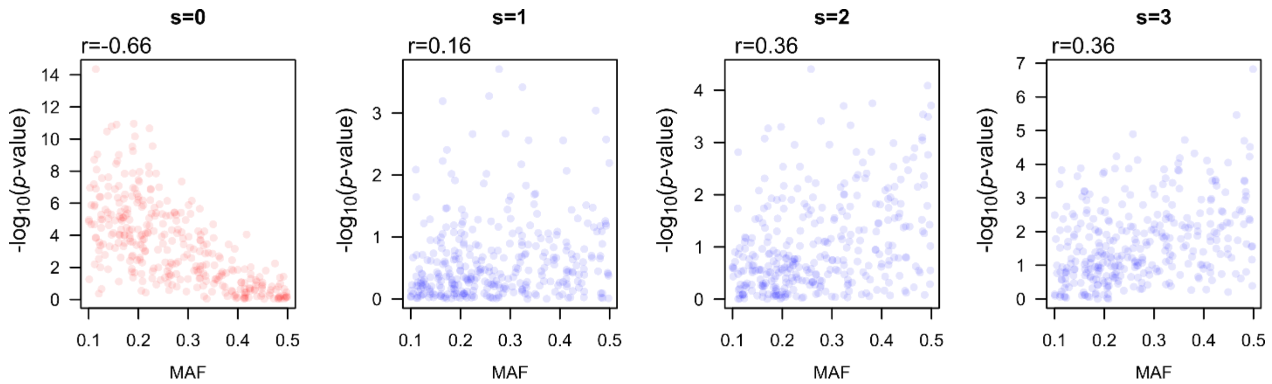


**Figure S9.** Relationship between the minor allele frequency (MAF) and false/true detection of the neighbor signals ($\beta_2 \neq 0$), when the collinearity and neighbor effects are both strong. Results among the ten iterations at total PVE = 0.8, $\alpha = 0.01$, $PVE_{self}$:$PVE_{nei}$ = 1:8, and the number of causal SNPs = 50, are shown. The scores of -$\log_{10}$($p$-values) for self ($s$ = 0) or neighbor ($s$ = 1 - 3) effects are plotted against the MAF. Pearson's correlation coefficients $r$ are listed above each panel.

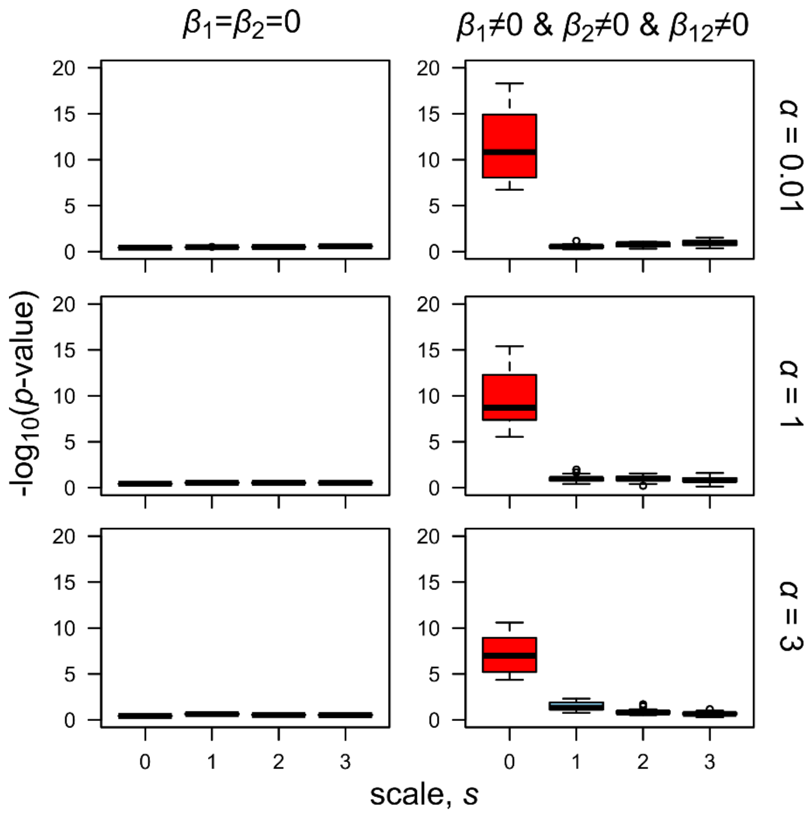


**Figure S10.** Association scores of the self-genotype effects (red) and symmetric neighbor effects (blue), when the asymmetric neighbor effects are strong. The score of $-\log_{10}$($p$-value) is averaged within each iteration, and shown for non-causal SNPs ($\beta_1 = \beta_2 = 0$), SNPs responsible for asymmetric neighbor effects ($\beta_1 \neq 0$ and $\beta_2 \neq 0$ and $\beta_{12} \neq 0$). The contribution of the asymmetric neighbor effects was set as large as $PVE_{self}$:$PVE_{nei}$:$PVE_{s\times n}$ = 1:1:8.

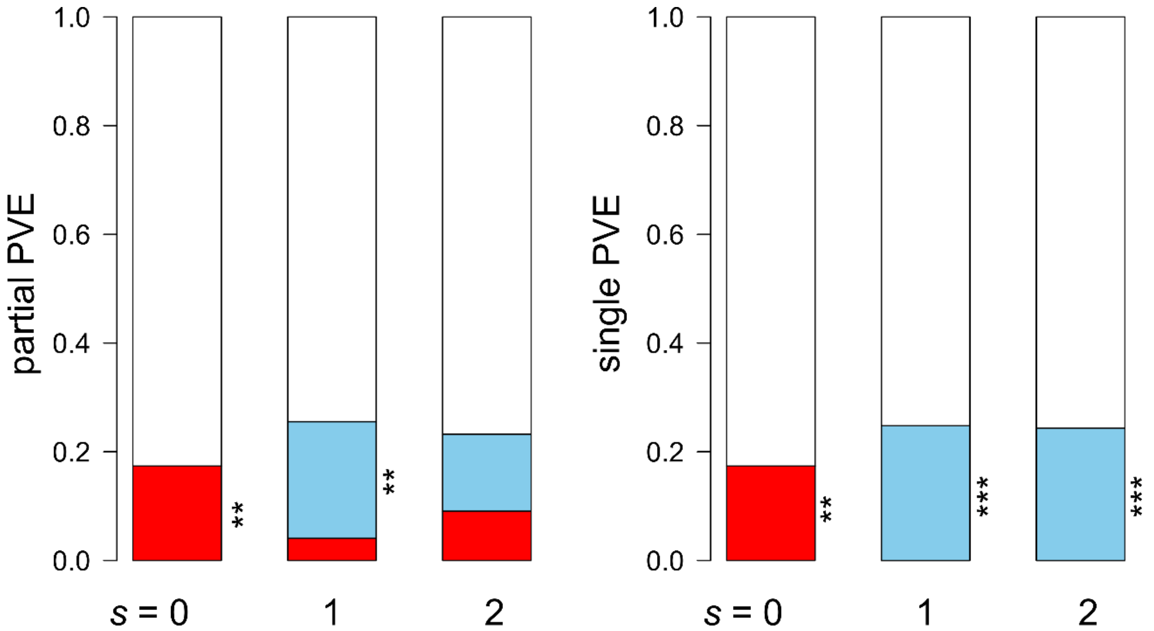


**Figure S11.** Proportion of the phenotypic variation explained (PVE) by the self-genotype or neighbor effects on the leaf damage score of field-grown *Arabidopsis thaliana*. The partial or single PVE is shown at the spatial scale of $s = 0$ (without any neighbor effects), $s = 1$ and $s = 2$, as defined by the partial PVE$_{\text{self}}$ $= \sigma_1^2/(\sigma_1^2 + \sigma_2^2 + \sigma_e^2)$; partial PVE$_{\text{nei}}$ $= \sigma_2^2/(\sigma_1^2 + \sigma_2^2 + \sigma_e^2)$; single PVE$_{\text{self}}$ $= \sigma_1^2/(\sigma_1^2 + \sigma_e^2)$; and single PVE$_{\text{nei}}$ $= \sigma_2^2/(\sigma_2^2 + \sigma_e^2)$. The red or blue fraction indicates PVE by the self or neighbor effects, respectively. The white fraction indicates residuals. Asterisks highlight a significant fraction with stepwise likelihood ratio tests from simpler to complex models: $^{**}p$-value < 0.01: $^{***}p$-value < 0.001.

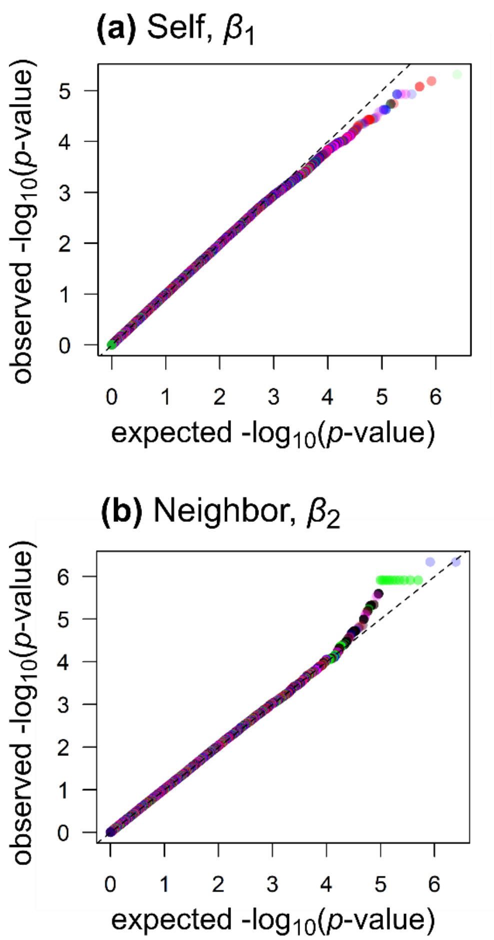


**Figure S12.** QQ-plots of the self and neighbor effects on the leaf damage score of *Arabidopsis thaliana*. Observed $-\log_{10}$($p$-values) scores are plotted against the expected ones. Dashed line indicates the same value between the observed and expected score. Plot colors represent the chromosome numbers as shown in the Manhattan plot (Fig. 6).

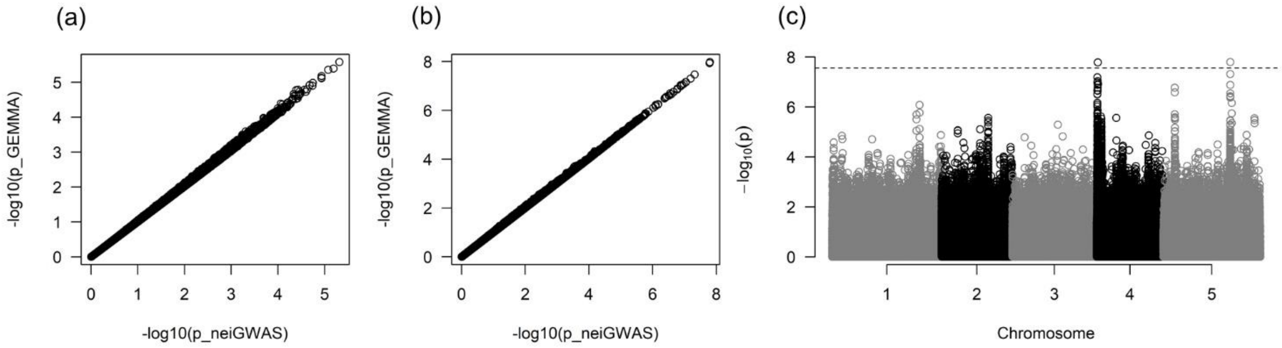


**Figure S13.** Comparison of self *p*-values between GEMMA and neighbor GWAS. The -$\log_{10}$(*p*-value) scores calculated by GEMMA or rNeighborGWAS are shown for the present leaf damage data (a) or publicly available flowering time data (b). Manhattan plot for the flowering time data (c) is depicted using *p*-values obtained by rNeighborGWAS, where a horizontal dashed line indicates a genome-wide Bonferroni threshold at $p = 0.05$.
